# Does powder averaging remove dispersion bias in diffusion MRI diameter estimates within real 3D axonal architectures?

**DOI:** 10.1101/2021.04.15.439974

**Authors:** Mariam Andersson, Marco Pizzolato, Hans Martin Kjer, Katrine Forum Skodborg, Henrik Lundell, Tim B. Dyrby

## Abstract

Noninvasive estimation of axon diameter with diffusion MRI holds potential to investigate the dynamic properties of the brain network and pathology of neurodegenerative diseases. Recent methods use powder averaging to account for complex white matter architectures, such as fibre crossing regions, but these have not been validated for real axonal geometries. Here, we present 120 – 313 *μ*m long segmented axons from X-ray nano-holotomography volumes of a splenium and crossing fibre region of a vervet monkey brain. We show that the axons in the complex crossing fibre region, which contains callosal, association, and corticospinal connections, are larger and exhibit a wider distribution than those of the splenium region. To accurately estimate the axon diameter in these regions, therefore, sensitivity to a wide range of diameters is required. We demonstrate how the *q*-value, *b*-value, signal-to-noise ratio and the assumed intra-axonal parallel diffusivity influence the range of measurable diameters with powder average approaches. Furthermore, we show how Gaussian distributed noise results in a wider range of measurable diameter at high *b*-values than Rician distributed noise, even at high signal-to-noise ratios of 100. The number of gradient directions is also shown to impose a lower bound on measurable diameter. Our results indicate that axon diameter estimation can be performed with only few *b*-shells, and that additional shells do not improve the accuracy of the estimate. Through Monte Carlo simulations of diffusion, we show that powder averaging techniques succeed in providing accurate estimates of axon diameter across a range of sequence parameters and diffusion times, even in complex white matter architectures. At sufficiently low *b*-values, the acquisition becomes sensitive to axonal microdispersion and the intra-axonal parallel diffusivity shows time dependency at both in vivo and ex vivo intrinsic diffusivities.

## Introduction

Action potentials propagate along axons and enable communication across different parts of the central nervous system. The axonal morphologies are crucial for the signal conduction process [1–3] and determine the conduction velocity (CV) with which signals are propagated. By electrophysiological modelling of the axon, Drakesmith et al. showed that axon diameter (AD) is the most important determinant of CV in myelinated axons [4]. AD is also a potential biomarker of neurodegenerative diseases such as Amyotrophic Lateral Sclerosis [5] and Multiple Sclerosis (MS) [6], and has been suggested to correlate with clinical scores of cognitive impairment in MS patients [7]. Thus, AD sheds light on brain health, as well as the structural and functional properties of the brain network.

Diffusion Magnetic Resonance Imaging (MRI) non-invasively probes the microstructural brain tissue environment by measuring the diffusion of water molecules across millisecond time scales. By fitting biophysical models that describe the underlying tissue microstructure to the diffusion MRI signal, AD can be estimated [8, 9]. The AxCaliber method [10, 11] uses pulsed gradient spin echo (PGSE) measurements with numerous combinations of gradient strengths and diffusion times to output the AD distribution (ADD), as demonstrated in vivo in the rat [10] and human [12] brains. AxCaliber requires prior knowledge of the axon orientation since it relies on measurements being made perpendicular to the axons. The ActiveAx approach has been demonstrated in vivo in humans and ex vivo in primates [13, 14] and outputs a mean AD index. Contrary to AxCaliber, ActiveAx is invariant to the orientation of the main fibre direction and implements an optimised acquisition consisting of three *b*–value shells, the minimum number of shells required to fit the three parameters in the signal model used by ActiveAx. These are sampled in ∼ 90 unique directions distributed uniformly on the unit sphere. These two methods have in common that they do not account for non-parallel axons, i.e. orientation dispersion (OD), or multiple bundles of crossing axons, factors which bias the AD measurement. Zhang et al. extended the ActiveAx approach in two ways. They relaxed the assumption of a single main fibre direction to enable AD estimation in regions of the ex vivo monkey brain in which there were crossing axon bundles [15], but the OD within those bundles was not taken into account. Later, Zhang et al. modelled the OD as a Watson distribution to fit AD in the in vivo human brain [16], but the method assumed a single main bundle direction. As such, diffusion MRI-based AD studies have mostly targeted the corpus callosum (CC), an organised white matter (WM) region that consists of aligned interhemispheric axonal connections. The CC has also been the subject of light and electron microscopy (EM) studies on AD, and these have been used as validation for the diffusion MRI-based AD metrics [10, 17, 18]. However, recent 3D imaging studies in the monkey and mouse CC demonstrate the complex morphologies, OD and trajectory variations of axons [19–21], and show how − even in the highly organised CC − these will bias AD measurements [19, 20, 22, 23]. Diffusion MRI-based estimates of AD should thus take into account three different classes of orientation effects: 1) the macroscopic fibre architecture, describing the relative orientations of different fibre bundles e.g. in crossing fibre regions; 2) the OD, describing the average dispersion exhibited by axons within each bundle; and 3) the microdispersion, describing the changes in trajectory and curvature along individual axons on the length scale of the measured diffusion.

The effects of the macroscopic fibre architecture and OD can be removed by powder averaging (PA). The PA involves calculating the arithmetic mean of the diffusion MRI signal in isotropically distributed directions on the unit sphere. Each diffusing spin can be described as probing a micro-domain, a microscopic region of the tissue environment within a voxel. The PA signal thus represents the spherical mean of the set of micro-domains, regardless of their individual orientation or organisation within the voxel. Several studies have used the PA to disentangle the effects of fibre architecture and OD from diffusion metrics [22, 24–33], and it has recently been implemented to estimate AD in the entire brain WM [34, 35]. To obtain estimates of AD index in the in vivo human brain, Fan et al. [34] fitted a multi-compartment spherical mean technique (SMT) model to the PA signal. The signal was sampled in up to 64 uniformly distributed directions for two diffusion times. In total, 16 unique *b*–values up to *b* =∼ 20 ms *μ*m^-2^ were acquired, enabled by the high in vivo gradient strengths of up to 300 mT/m of the Connectom scanner [36–38]. Veraart et al. [35], on the other hand, modelled only the intra-axonal space (IAS) by fitting a power law (PL) to the PA signal at high *b*–values that suppress the signal from the extra-axonal space (EAS) [39, 40]. This used b ≥ 20 ms *μ*m^-2^ for ex vivo experiments and *b* ≥ 6 ms *μ*m^-2^ for in vivo experiments. The signal was measured along 60 uniformly distributed gradient directions at a single diffusion time for up to 18 *b*-values. AD was calculated for the ex vivo rat brain and, also using a Connectom scanner, for the in vivo human brain.

Although the PA techniques remove fibre architecture and OD effects, they rely on the assumption that the micro-domain probed by diffusing spins is cylindrical. With increasing diffusion times, the spins diffuse further and increasingly probe the microdispersion of the axons, violating the assumption of a cylindrical micro-domain and making the AD estimate time-dependent. The effects of diameter and trajectory variations on estimated AD have been described [22, 41, 42], but the diffusion times and *b*-values for which the PA-based AD estimate becomes sensitive to the microdispersion in axons is unknown. The signal-to-noise ratio (SNR) of the signal [43] and the gradient strength of the applied magnetic field [14] affect the sensitivity profile of the acquisition, placing limits on the upper and lower bounds of measurable AD. How different sequence parameters, the number of gradient directions or the SNRs affect these bounds has not been investigated for PA-based AD estimates.

Validating the estimated ADs from the PA methods for different sequence parameters and in real axons is therefore important. The PL implementation from Veraart et al. has been evaluated within axon segments of length ∼ 20 *μ*m from electron microscopy (EM) of the mouse CC using Monte Carlo (MC) simulations of diffusion [22]. Given that non-axonal structures in the EAS can impact the trajectories of axons for up to 20 *μ*m [19], longer axonal segments may to a greater extent represent the characteristics of the IAS. Notably, although the PA is expected to factor out the effects of fibre crossings and OD, it has only been validated on segments of axons from the CC in which the fibre architecture is simple and does not contain substantial crossings.

In this study, we adopt a simulation-based modus operandi to validate PA-estimates of AD in real axons between 120 and 313 *μ*m in length, segmented from large field of view (FOV) X-ray nano-holotomography (XNH) volumes of two inherently different WM architectures of the vervet monkey brain. The two regions are: a) the ordered splenium CC (Fig. 1C) and b) a heterogeneous crossing fibre region (Fig. 1B). Throughout, we restrict the analysis to the IAS to analyse how accurately it is represented by the PA model at different *b*-values, and the results are relevant to both multi-compartment and single-compartment models that include the IAS. Firstly, we explore how the SNR, gradient strength (hence, also *b*-value) and number of unique gradient directions determine the sensitivity profile of the PA AD estimation method to different cylinder diameters. Secondly, we demonstrate the impact (or lack thereof) on estimated AD of using inaccurate assumptions of the parallel diffusivity for different *b*-values. Lastly, we validate the AD estimates from the SMT and PL at different diffusion times, gradient strengths (typical of pre-clinical or human Connectom scanners), diffusivities (in vivo/ex vivo) and, pertinently, within the complex segmented axons of the primate brain where ADs are similar to those of the human brain [44].

**Fig. 1:**
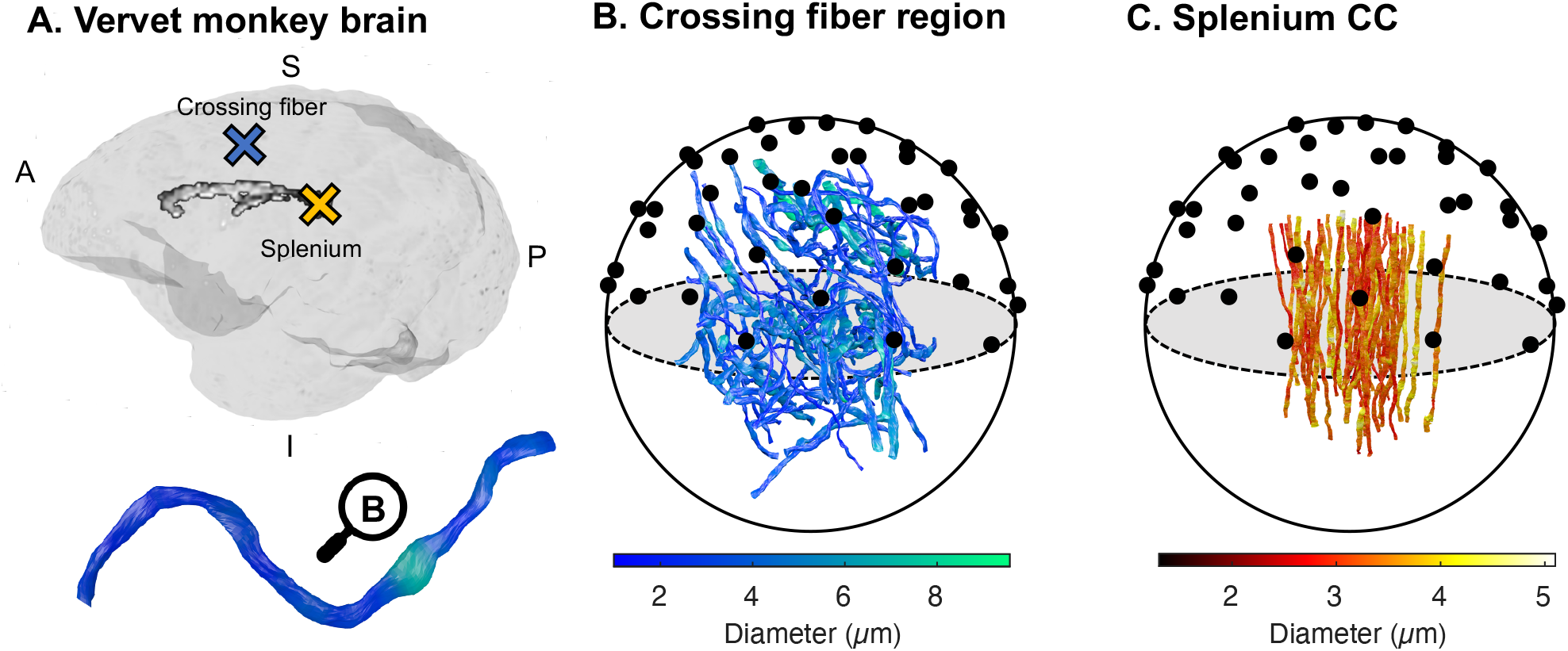
Can axon diameter be accurately estimated in different WM regions using PA approaches, despite complex axonal morphologies? Validation of PA-estimates of axon diameter was carried out in two different fibre architectures of the vervet monkey brain. A) Axons were segmented from XNH volumes of a heterogeneous crossing fibre region in the anterior semiovale (blue cross) and the splenium (yellow cross). B) 59 segmented crossing fibre axons and C) 58 segmented splenium axons from [19]. The axon diameters are indicated by the respective colorbars and the black circles represent uniformly distributed directions on the unit sphere.

## Theory

To model the IAS, the SMT and PL approaches both assume that the micro-domains probed by spins within the IAS are cylindrical. Here, we present an outline of the origins of the SMT expression for cylinders, based on the theory presented elsewhere [45, 46]. It is this expression that is used to represent the IAS in the SMT-based approach used in Fan et al. [34]. From the SMT expression, the assumption of high *b*-values entails that the SMT can be formulated as a PL, as that in Veraart et al. [35].

### Modelling the powder averaged signal in a cylinder

In a cylinder, the apparent diffusion coefficient (ADC) at any angle *α* to its axis is [47]:

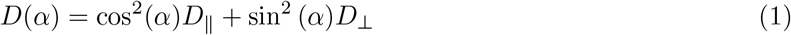

where *D*_‖_ is the diffusivity parallel to the cylinder axis and *D*_⊥_ is that perpendicular to it, and carries the information regarding cylinder diameter. Powder averaging of the diffusion MRI signal involves integrating it over an infinite number of uniformly distributed inclination angles *α* relative to the cylinder axis to give the powder averaged signal, 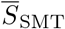 [45] :

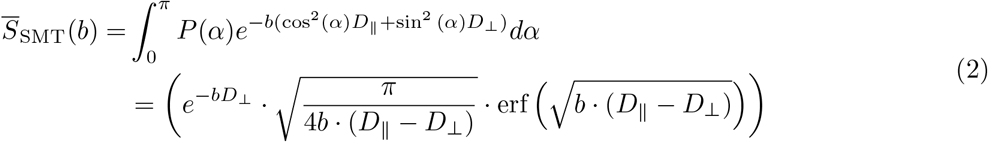

where 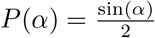 ensures that the weighting of the angles is uniform, erf(*x*) is the error function of *x*, and *b* is the diffusion weighting. Eq. 2 is what is here referred to as the “SMT implementation” and is the analytical description of the spherically averaged signal within a cylinder. Aside from the *b*-value, the signal depends on two variables: *D*_‖_ and *D*_⊥_. Only a finite number of directions, *N*, can be used in practice. As such, *N* is one of the variables that determines the accuracy of the measurement.

In cylinders, where *D*_‖_ > *D*_⊥_ and at high *b*-values, it can be assumed that *b* · (*D*_‖_ – *D*_⊥_) ≫1. In these conditions, erf(*x*) = 1 and Eq. 2 can be rearranged to take the form of a PL:

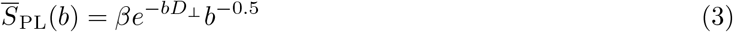

where 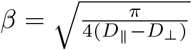. Eq. 3 is what is here referred to as the “PL implementation” and is an alternative representation of the SMT at high *b*–values.

In practise, whether using single- or multi-compartment models of the WM, the fraction of the total signal that the IAS represents, *f*_*a*_, is unknown. It thus needs to be included as a multiplicative constant in Eqs. 2 or 3. This introduces an additional third variable into the SMT implementation in Eq. 2, such that the signal depends on *f*_*a*_, *D*_‖_ and *D*_⊥_. In the case of the PL, *f*_*a*_ can simply be incorporated into the existing constant *β* such that 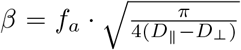. This removes the need to fit a third parameter to the PA signal.

### Converting *D*_⊥_ into a diameter

By fitting Eqs. 2 or 3 to the PA signal, the ADC perpendicular to the cylindrical micro-domains, *D*_⊥_, is obtained. The cylinder diameter can be calculated from *D*_⊥_ using the formulation in van Gelderen et al. [47]:

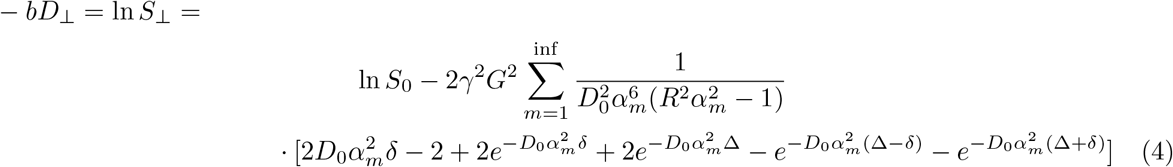

where *S*_⊥_ is the diffusion weighted signal perpendicular to the cylinder, *S*_0_ is the signal with no diffusion weighting, *γ* is the gyromagnetic ratio, *G* is the strength of the gradient pulse, *D*_0_ is the intrinsic diffusivity, *δ* is the duration of the gradient pulse, Δ is the separation of the gradient pulses, *R* is the cylinder radius and *α*_*m*_ is the mth root of 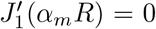 where 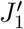 is the derivative of the first order Bessel function of the first kind. In this study, Eq. 4 is calculated up to *m* = 6.

If 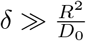, as can be assumed to be the case for most axons [35], the cylinders are said to fall within the Neuman limit [48, 49] and Eq. 4 simplifies to:

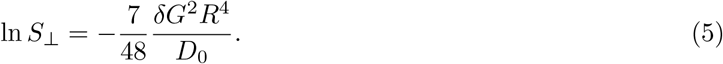

Importantly, both the SMT and PL require knowledge – or an assumption – of *D*_0_ in order to estimate a diameter from *D*_⊥_. However, since 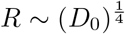 in Eq. 5, R is relatively insensitive to small inaccuracies of *D*_0_.

## Materials and Methods

### Simulations

The simulations in this study are divided into two categories: simulations of the diffusion MRI signal from cylinders of different diameters and simulations in the IAS of segmented axons from XNH volumes of the vervet monkey brain presented in [19]. By simulating the signal from cylinders, the impact of scanning parameters, SNR and model assumptions on estimated diameter could be isolated. By simulating diffusion within segmented axons from the splenium and crossing fiber regions, the impact on the estimated AD of real fiber architectures, OD and microdispersion was investigated.

### Simulating the diffusion MRI signal from cylinders

The signals arising from cylinders of different diameter, aligned with the *z*-axis, were generated analytically. For given PGSE parameters *δ*, Δ and *G* and radius *R*, Eq. 4 was used to calculate *D*_⊥_ for each cylinder. From this, the ADC and signal in any normalised gradient direction *ê*_*G*_ = [*e*_*x*_,*e*_*y*_, *e*_*z*_] could be calculated from 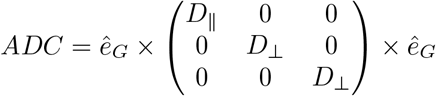.

### Simulating diffusion within the real IAS from XNH volumes of the monkey brain

We used segmented axons from the brain of a 32-month old female vervet monkey, imaged with 3D synchrotron XNH acquired at the European Synchrotron Research Facility, beamline ID16A. The axons originated from two different brain regions: the splenium of the CC and a “crossing fiber region”, located in a position of the anterior centrum semiovale where the diffusion MRI data indicated the crossing of the corticospinal tract, interhemispheric callosal fibers and association fibers [19]. A description of both XNH volumes, as well as the segmentation and analysis of splenium axons is given in Andersson et al [19]. In short, the XNH volume of the splenium had an isotropic voxel size of 75 nm and cylindrical FOV of diameter and length 153.6 *μ*m. From the splenium, 54 axons of minimum length 120 *μ*m were segmented at the native 75 nm image resolution. The XNH volume of the crossing fiber region had an isotropic voxel size of 100 nm and cylindrical FOV of diameter and length 204.8 *μ*m. The much larger diameters of axons in this region entailed that the segmentation of 58 axons of minimum length 120 *μ*m could be manually performed in ITK-SNAP [50] (RRID:SCR_002010) at a downsampled isotropic voxel size of 500 nm. This was significantly less time consuming than segmenting the axons at higher resolution. Smaller axons were present in both XNH volumes, but could not be segmented due to their small diameters in comparison to the voxel size and low SNR [19]. After segmentation, the equivalent diameters of the axons were quantified in the plane perpendicular to their local trajectory [19]. The volume-weighted AD, *d*, of each axon was estimated as 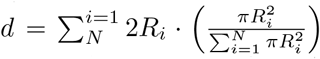 where *R*_*i*_ is the ith measured radius of *N* equidistant measurement points along the axonal trajectories. The volume-weighted mean diameters of the population of splenium axons and crossing fiber axons were similarly calculated.

The axon segmentations were converted to triangulated surface meshes, after which the Monte Carlo Diffusion and Collision (MCDC) framework [51] was used to simulate diffusion within each axon mesh. The simulations used an intrinsic ex vivo diffusivity of *D*_0_ = 0.6 *μ*m^2^ms^-1^, 2 · 10^5^ uniformly distributed spins per axon and 1 · 10^−5^ seconds per time step, as in Andersson et al [19]. Simulations were also performed using an in vivo diffusivity of *D*_0_ = 2 *μ*m^2^ms^-1^ to mimic diffusion within the living human brain, but with 3.4 · 10^−6^ seconds per time step to ensure the same step length at the higher diffusivity. Initialisation of the spins was performed at a minimum distance from the ends of the axons (minimu 20 *μ*m for ex vivo simulations and 30 *μ*m for in vivo diffusivities), to prevent their escape from the IAS with regards to the diffusivity and the maximum diffusion times used (∼ 40 ms for ex vivo simulations and ∼ 30 ms for in vivo simulations).

### Diffusion MRI scanning parameters

For all experiments, the PGSE waveform was used. Throughout the investigation, different sequence parameters were varied to isolate the effects of different variables on the estimated diameter.

In the simulations on cylinders, a gradient duration of *δ* = 7.1 ms was used, similar to in [34, 35]. The gradient separation was kept at Δ = 20 ms, and the effective diffusion time, *t*_*d*_, was given by 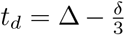 The simulations on cylinders used only ex vivo diffusivities, and *b*-values were referred to as ‘‘high” if they surpassed ≳20 ms *μ*m^-2^, the value at which the EAS was said to be suppressed in [35]. Most simulations in cylinders used high *b*-values in the range of *b* = [19.25, 63.62] ms *μ*m^-2^ to allow a direct comparison between the SMT and PL implementations, the latter of which required the use of high b-values. Different *b*-values were obtained by varying the gradient strength, *G*, given that *b* = *q*^2^*t*_*d*_ where *q* is the diffusion encoding *q* = *γδG* [m^-1^].

In the simulations of the real IAS, *δ* = 7 ms was used. Both *G* and Δ were varied to assess the effects of different diffusion times. In these simulations, the SMT and PL were fitted to many different *b*-values, ranging between *b* = [0.55, 65] ms *μ*m^-2^ for ex vivo diffusivities and *b* = [0.44, 8.73] ms *μ*m^-2^ for in vivo diffusivities.

For the most part, fits of the SMT and PL to the PA signal from several *b*-values used three shells, similar to ActiveAx [13, 14], since three was the minimum number of shells needed to calculate *f*_*a*_, *D*_‖_ and *D*_⊥_ in the SMT implementation. Uniformly distributed directions on the unit sphere were generated according to the electrostatic repulsion method [52, 53].

### Distinguishing the signal from noise

The effect of noise on the AD estimation was studied by adding Rician noise of variable SNR to the noise-free, normalised signals. The total variance of the noise was defined as 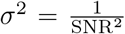. Rician distributed noise was simulated calculating the magnitude of complex Gaussian noise in which the real and imaginary components each had a standard deviation of 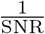 [54].

To assess whether or not a single signal could be distinguished from noise at a given SNR and a single *b*-value, we used the sensitivity criterion of Nilsson et al. [43] for parallel cylinders. The smallest robustly measurable change of the normalised signal, Δ*S* was defined as:

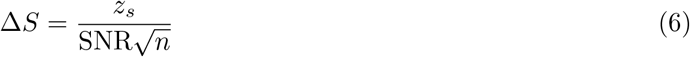

where *n* was the number of repeated measurements and *z*_*s*_ was the z-threshold for the significance level s. The signal was thus said to be sensitive between the bounds [Δ*S*, 1 – Δ*S*]. The diameters that gave rise to the PA signal at these boundaries were defined as the maximum and minimum bounds of the measurable diameter. Here, we choose s = 0.05, giving *z*_*s*_ = 1.64, as in [43].

To predict whether the PA signal could be distinguished from normally distributed noise, the sensitivity criterion of Nilsson et al. [43] for fully dispersed cylinders was used. It is defined as:

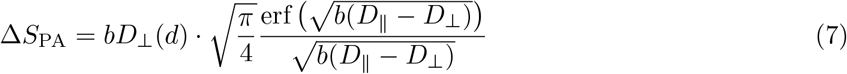

where *D*_⊥_ (*d*) is the perpendicular diffusivity of the diameter, *d*, that is defined as:

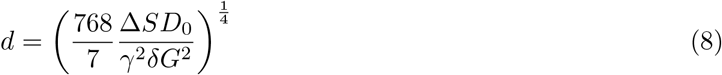

When using Eqs. 7 and 8, *n* in Eq. 6 was set to the number of unique gradient directions. From Δ*S*_PA_, the theoretical range of measurable diameters was calculated as the diameters with PA signals within the range [Δ*S*_PA_, *S*_stick_ – Δ*S*] where *S*_stick_ is the signal of a cylinder with diameter equal to zero:

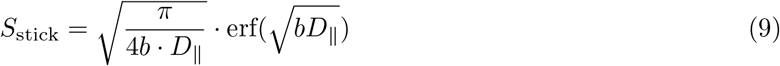

Importantly, Eqs. 6-8 are formulated for single *b*-values only and assume that the noise follows a normal distribution.

### Fitting the Spherical Mean Technique and Power Law to the PA signal

#### The Spherical Mean Technique implementation

The signal fraction of the IAS, *f*_*a*_, was incorporated into the SMT formulation in Eq. 2 such that:

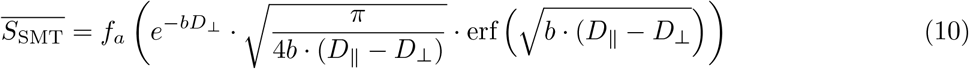

The SMT was fitted to the PA signals with the non-linear least squares solver `lsqnonlin()` in Matlab R2020a. Once *D*_⊥_ had been fitted, the diameter was calculated using Eq. 4. We implemented three different variations of the SMT fit to assess the robustness when keeping different variables fixed:

i. **SMT-1**: a single-shell fit to obtain *D*_⊥_ in the range [0, *D*_‖_]. Assumes known *f*_*a*_ and ex vivo *D*_⊥_ = 0.6 *μ*m^2^ ms^-1^.
ii. **SMT-2**: a multi-shell fit to obtain *D*_⊥_ in the range [0, *D*_‖_] and *f*_*a*_ in the range [0, 1]. Assumes known ex vivo *D*_‖_ = 0.6 *μ*m^2^ms^-1^, or in vivo *D*_‖_ = 2 *μ*m^2^ms^-1^.
iii. **SMT-3**: a multi-shell fit to obtain *D* _⊥_ in the range [0, *D*_0_ · 1.5], *f*_*a*_ in the range [0,1] and *D*_‖_ in the range[D_0_/2, *D*_0_ · 1.5] where *D*_0_ was the known intrinsic diffusivity of the simulations. For ex vivo and in vivo simulations, *D*_0_ = 0.6 *μ*m^2^ms^-1^ and *D*_0_ = 2 *μ*m^2^ms^-1^ were used respectively.

SMT-1 was used to assess the effect of the number of directions and SNR on the estimated diameter in the best case scenario in which *f*_*a*_ and *D*_‖_ are known. Moving to a more realistic scenario, SMT-2 was used to assess the accuracy of SMT-based diameter estimation at different, unknown values of *f*_*a*_. SMT-2 was also used to investigate the consequences of enforcing an incorrect value of *D*_‖_ at different b–values and SNRs. Lastly, SMT-3 placed no assumptions on any of the variables, similarly to the PL implementation. SMT-3 was used to investigate the ability of the SMT to estimate AD in real axons for in vivo and ex vivo intrinsic *D*_0_, assuming no prior knowledge of diffusivities (other than their upper and lower bounds). SMT-3 also outputted estimates of *D*_‖_ in the axons.

#### The Power Law implementation

To assess the diameter estimates from the PL formulation, the expression in Eq. 3 was fitted to the PA signal from cylinders of different diameters, providing estimates of *D*_⊥_ and *β* This allowed for a comparison of the PL-derived diameter, *d*_PL_, with those from SMT-2 and SMT-3. To fit the PL the Matlab-based nonlinear least squares estimator provided by Veraart and Novikov [55] was used. The implementation assumed the Neuman limit in Eq. 5 to obtain a diameter estimate, as in Veraart et al. [35].

## Results

The Results section is organised as follows. First, we perform an analysis of the range of diameters to which the PA maintains sensitivity for different SNRs and gradient strengths to verify that it would theoretically be possible to calculate the diameters of the XNH-segmented axons. We proceed to verify the multi-shell SMT-2 and PL descriptions for different numbers of shells, and investigate the effects of both Rican and Gaussian distributed noise on the diameter estimates. The SMT-2 and PL formulations implement different assumptions and degrees of a priori knowledge. We therefore evaluate whether the assumption of a known parallel diffusivity is valid, using both low and high *b*-values. Then, we present segmentations of axons from the splenium and a crossing fiber region of the monkey brain. In these, we investigate the ability of the SMT-2, SMT-3 and PL implementations to estimate the volume-weighted ADs within real axonal geometries for in/ex vivo diffusivities and different *b*-values, given by combinations of different gradient strengths and diffusion times between ∼ 10 – 40 ms.

### Angular sensitivity of the diffusion MRI signal in cylinders

Diameter estimates using the PA demand that the diffusion MRI signal is sensitive to the different length scales probed by measuring the signal at inclination angles *α* relative to the cylinder axis. The range of diameters that can be measured with the PA can be predicted from Eq. 7, and is shown in Fig. 2B for a range of SNRs and *q*-values. The widening/narrowing of the sensitivity range can be explained by the sensitivity of the PGSE acquisition to different angles *α*. As shown in Fig. 2C, the range of a to which there was sensitivity varied with the SNR and the diffusion encoding *q* (and hence the gradient strength). The angular sensitivity profiles also varied according to the diameter of the cylinders. With increasing SNR (Fig. 2C, top row), the sensitivity to both the high and low *α* increased. At SNR = 100, an increasing *q* increased the sensitivity to high *α*, but the additional attenuation of the signal caused by the higher diffusion weighting decreased the sensitivity to small *α*.

**Fig. 2:**
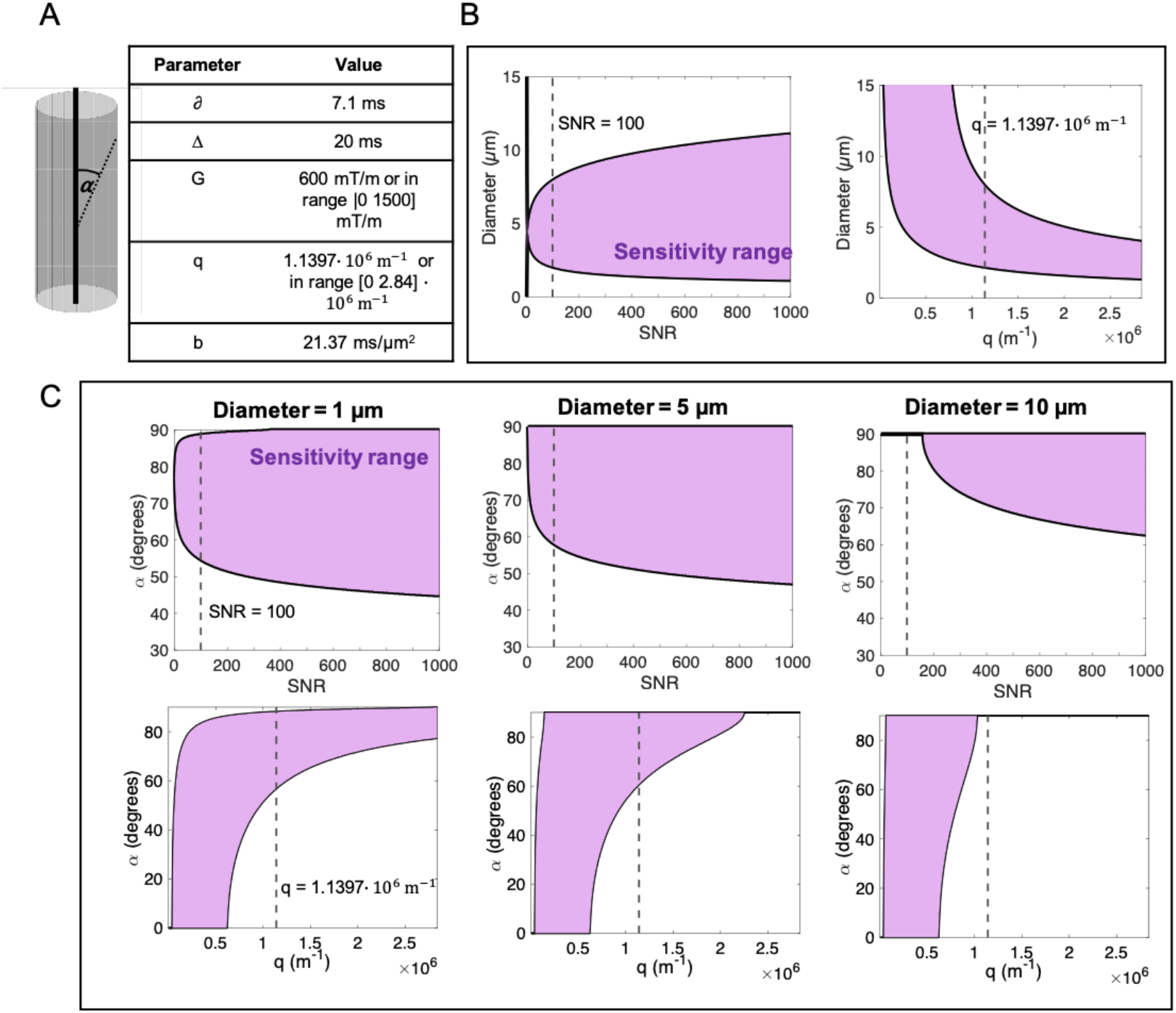
Angular sensitivity of the signal with respect to the cylinder axis. A) The angle *α* is defined as the inclination from the cylinder axis. The PGSE parameters in the table were used for the sensitivity analysis. B) The range of measurable diameters using the PA signal varies with the SNR, as shown using *q* = 1.1397 · 10^6^ m^-1^. For SNR = 100, the range of measurable diameters varies with the *q*-value (*G* is varied to obtain different *q*, but *δ* and Δ are as in the table). The sensitivity analysis is based on Eq. 7 and assumes 30 gradient directions. C) Variation with SNR and *q*-value of the angular sensitivity range, in terms of *α*, to which the measurement is sensitive in cylinders of diameter [1, 5, 10] *μ*m for the PGSE parameters in B. This sensitivity criterion is as in Eq. 6.

### Number of gradient directions

The number of gradient directions determines how well the PA signal represents the analytical PA of a cylinder. Although it would be ideal to use very many directions, a compromise factoring in limited scan time must be made. To investigate how the angular resolution affects the diameter estimate, the signals from cylinders of different diameters with different numbers of gradient directions (6, 15, 30 and 512) and levels of Rician noise were generated, emulating the noise distribution present in magnitude diffusion MRI images. The sequence parameters were the same as in Fig. 2 and SMT-1 was fitted to the PA signal, giving a diameter estimate *d*_SMT-1_, as shown in Fig. S1. The diameter from the signal measured in the one direction perpendicular to the cylinders (using Eq. 4), *d*_VG_, was also calculated. The theoretical upper and lower bounds of *d*_SMT-1_ were calculated from Eq. 7, while those of *d*_VG_ were calculated from Eq. 6.

We found that the number of gradient directions imposed an additional lower bound of measurable diameter separate from that incurred by finite SNR. Of the number of directions investigated, the prediction of the upper and lower bounds using Eq. 7, which assumes an infinite number of directions, was therefore only accurate starting from 30 directions using the given sequence parameters. At SNR= 20, there was a general underestimation of diameter regardless of the number of gradient directions, as a result of the Rician bias [54]. This disappeared when Gaussian distributed noise was used, as shown in Fig. S3. Furthermore, we noted a dependence of *d*_SMT-1_ on the orientation of the cylinder for fewer gradient directions than 30, as discussed in Supplementary Information S1 and shown in Fig. S2.

### Estimating diameter and intra-axonal signal fraction from multiple high *b*–value shells

The impact of different high *b*-values on diameter estimation was studied by fitting SMT-2 and the PL to the PA signals from cylinders with ground truth values of *f*_*a*_ = 0.8, an approximation of the expected axonal signal fraction. The *b*-values were chosen to cover a similar range to those in the ex vivo acquisition in [35], and all were ≳20 ms *μ*m^-2^ to simulate suppression of the EAS. For one set of ten b-values, Fig. 3 shows the estimated parameters from the SMT-1, SMT-2 and PL fits at SNR = 100 with Rician and Gaussian distributed noise.

**Fig. 3:**
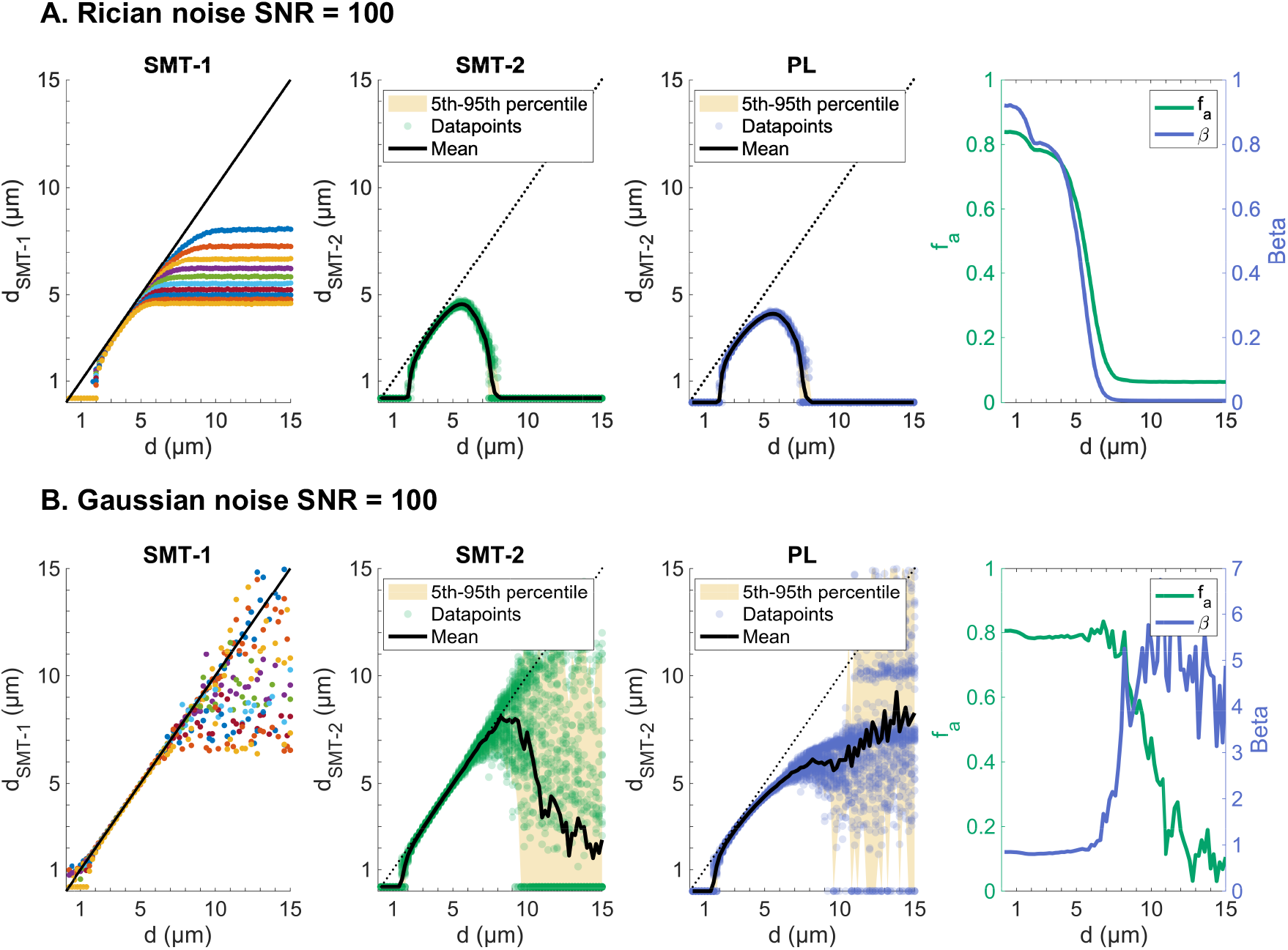
Multi-shell fit of SMT and PL to the signal from cylinders of diameter between 0.2 and 15.0 μm at SNR = 100 and different *f*_*a*_. Fitted *d*_SMT-1_, *d*_SMT-2_, *d*_PL_, *f*_*a*_ and *β* for ground truth A) *f*_*a*_ = 1 B) *f*_*a*_ = 0.8 and C) *f*_*a*_ = 0.5. The signal is generated using *D*_‖_ = 0.6 *μ*m^2^ms^-1^, 30 isotropically distribued directions and PGSE parameters *δ* = 7.1 ms, Δ = 20 ms, *G* = [550, 750, 1000] mT/m, *b* = [19.25, 35.79, 63.62] ms *μ*m^-2^. All datapoints reflect the mean of *N* = 50 measurements, and the error bars reflect their standard deviation.

For Rician noise in Fig. 3A, *d*_SMT-2_ and *d*_PL_ dropped to 0 at small and large diameters, in contrast to the single shell SMT-1 which plateaued at large diameters (Figs. S1, S3). At large diameters, the underestimation of diameter was accompanied by a reduction of estimated *f*_*a*_ for the SMT-2 approach as the significant attenuation at large diameters was instead attributed to a smaller signal fraction. A reduction in *β* was seen for the PL approach. The use of data with Gaussian distributed noise in Fig. 3B produced *d*_SMT-2_, *f*_*a*_ and *β* estimates that were accurate at smaller and larger diameters than with Rician noise, albeit with a high variance at large diameters. Consequently, at diameters ≳ *8 μm*, the estimates of *f*_*a*_ and *β* could not be robustly estimated, even with the mean of n = 50 repeats. One contributor to the high variance was that the SMT-2 and PL fits failed for many large diameters, defaulting to 0.

To explore how the *b*-value range and number of shells affected the diameter estimates, the experiments in Fig. 3 were repeated for A) three shells that spanned the same *b*-value range range with *b* = [19.25, 35.79, 63.62] ms *μ*m^-2^, B) three closely spaced *b*-values at the lower end of the range *b* = [19.25, 22.90, 26.88] ms *μ*m^-2^ and C) three closely spaced *b*-values at the higher end of the range, *b* = [51.54, 57.42, 63.62] ms *μ*m^-2^. The results are shown in Fig. 4. Other than a small decrease in variance (likely due to the increased number of sampling points), there seemed to be no clear advantage to sampling more shells that cover the same range of *b*-values, as clear from the comparison of the results using ten shells in Fig. 3A and using three shells in Fig. 4A. Secondly, fitting to the three *b*-values on the lower end of the range in Fig. 4B resulted in a wider range of measurable diameters than fitting to three significantly higher *b*-values in Fig. 4C. The equivalent experiments using Gaussian noise are shown in Fig. S4. Here, the use of Gaussian noise widened the range of measurable diameters and provided more accurate estimates of *f*_*a*_.

**Fig. 4:**
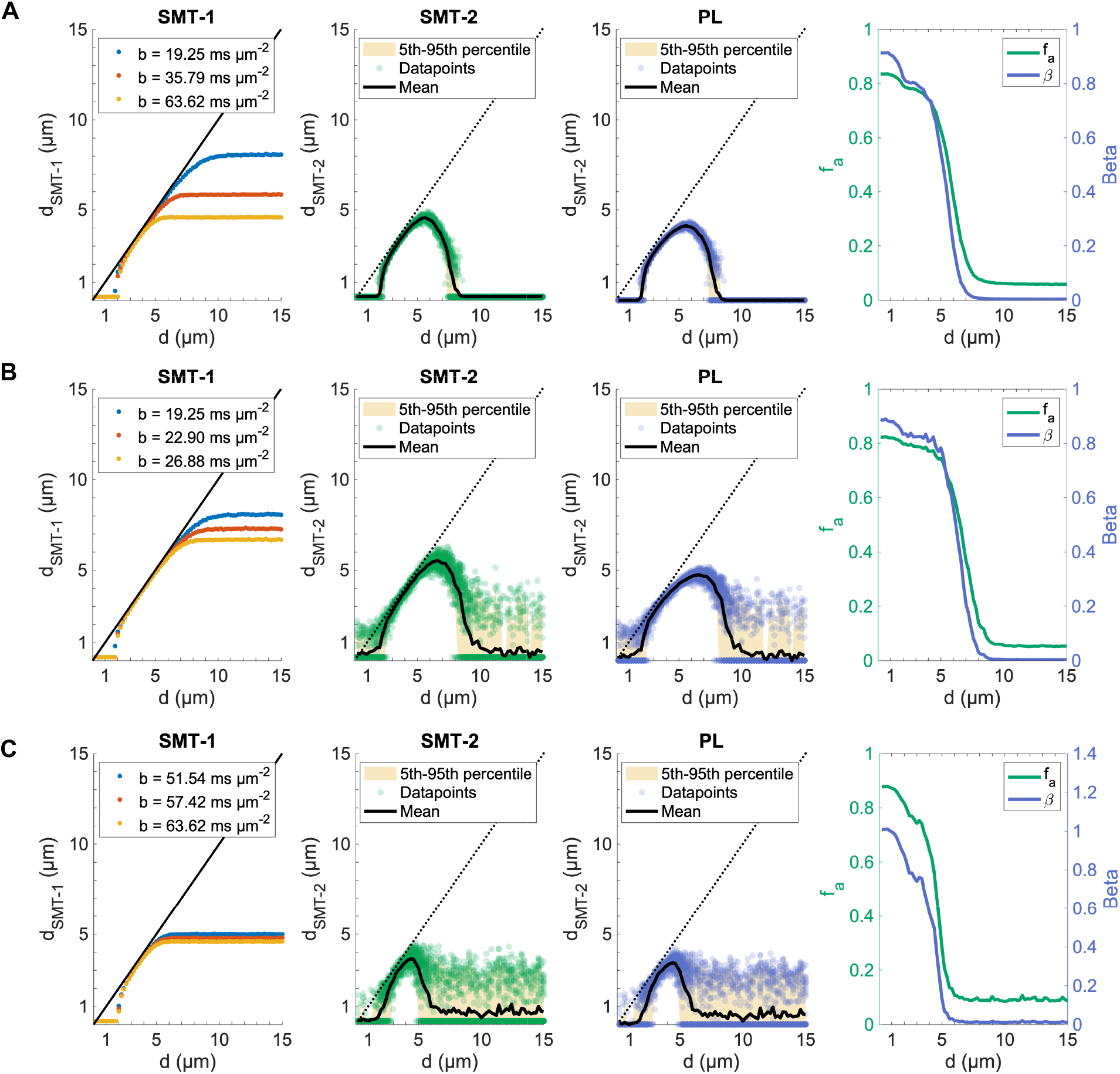
The choice of *b*-values affects the range of measurable diameters. Multi-shell fit of SMT and PL to the signal from cylinders of diameters between 0.2 and 15.0 *μ*m at SNR = 100 (Rician noise) and *f*_*a*_ *=* 0.8. Fitted *d*_SMT-1_, *d*_SMT-2_, *d*_PL_, *f*_*a*_ and *β* for ground truth A) three shells with *b =* [19.25, 35.79, 63.62] B) three shells with *b* = [19.25, 22.90, 26.88] ms *μ*m^-2^ and C) three shells with *b* = [51.54, 57.42, 63.62] ms *μ*m^-2^. The signal was generated using *D*_‖_*=* 0.6 *μ*m^2^ms^-1^, 30 directions, PGSE parameters *δ* = 7.1 ms, Δ = 20 ms and varying *G. n* = 50 repeats of each acquisition were performed for each diameter.

How different noise levels influenced the diameter estimates from the SMT-2 and PL fits was examined by repeating the experiment in Fig. 4A for SNR = [20, ∞], as shown in Fig. S5. We also examined whether the slight underestimation of diameter using the PL implementation contra the SMT-2 implementation in Figs. 3 and 4 could be due to the assumption of the Neuman limit (Eq. 5), as opposed to the full Gaussian phase approximation formulation from van Gelderen et al. [47] (Eq. 4) by calculating *d*_PL_ using both expressions. At SNR = ∞, *d*_SMT-2_ and *f*_*a*_ were accurately estimated up to ∼ 13 *μ*m, aside from at very small diameters. Importantly, we found that the underestimation seen in *d*_PL_ could indeed be explained 2 by the fact that large diameters *≥* 4 *μ*m could no longer be assumed to fulfil the assumption of 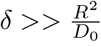 in the Neuman limit, and that using the full expression in Eq. 4 ensured equivalent accuracy of *d*_SMT-2_ and *d*_PL_ up to large diameters. As such, the underestimation of diameter in the PL implementation is not related to the PL signal model in Eq. 3. Rather, it is related to the conversion of *D*_‖_ into a diameter. As expected, reducing the SNR to 20 increased the variance of the estimates and decreased the range of measurable diameters. This effect was particularly prominent for Rician distributed noise, but also for Gaussian distributed noise where 50 repetitions were not sufficient to robustly estimate the diameter, *f*_*a*_ or *β* for diameters ≥ 6 *μ*m. Since *f*_*a*_ multiplies the signal attenuation, a change of its value directly determined a change in the SNR and the range of measurable diameters (Fig. S6).

### The dependence of estimated diameter on *D*_‖_

Whether or not SMT-2 can be fitted to a multi-shell acquisition depends on how robust the estimation of diameter, *d*_SMT-2_, is to inaccuracies in the assumed prior *D*_‖_ for the given diffusion time. For cylinders, the estimated *d*_SMT-2_ and *d*_*VG*_ for different assumed values of *D*_‖_ are shown in Fig. 5, with ground truth *D*_‖_ = 0.6 *μ*m^2^ms^-1^. For the diameter estimation, it was assumed that *D*_*0*_ = *D*_‖_ for all *D*_‖_ The range *D*_‖_ ± 15% is marked.

**Fig. 5:**
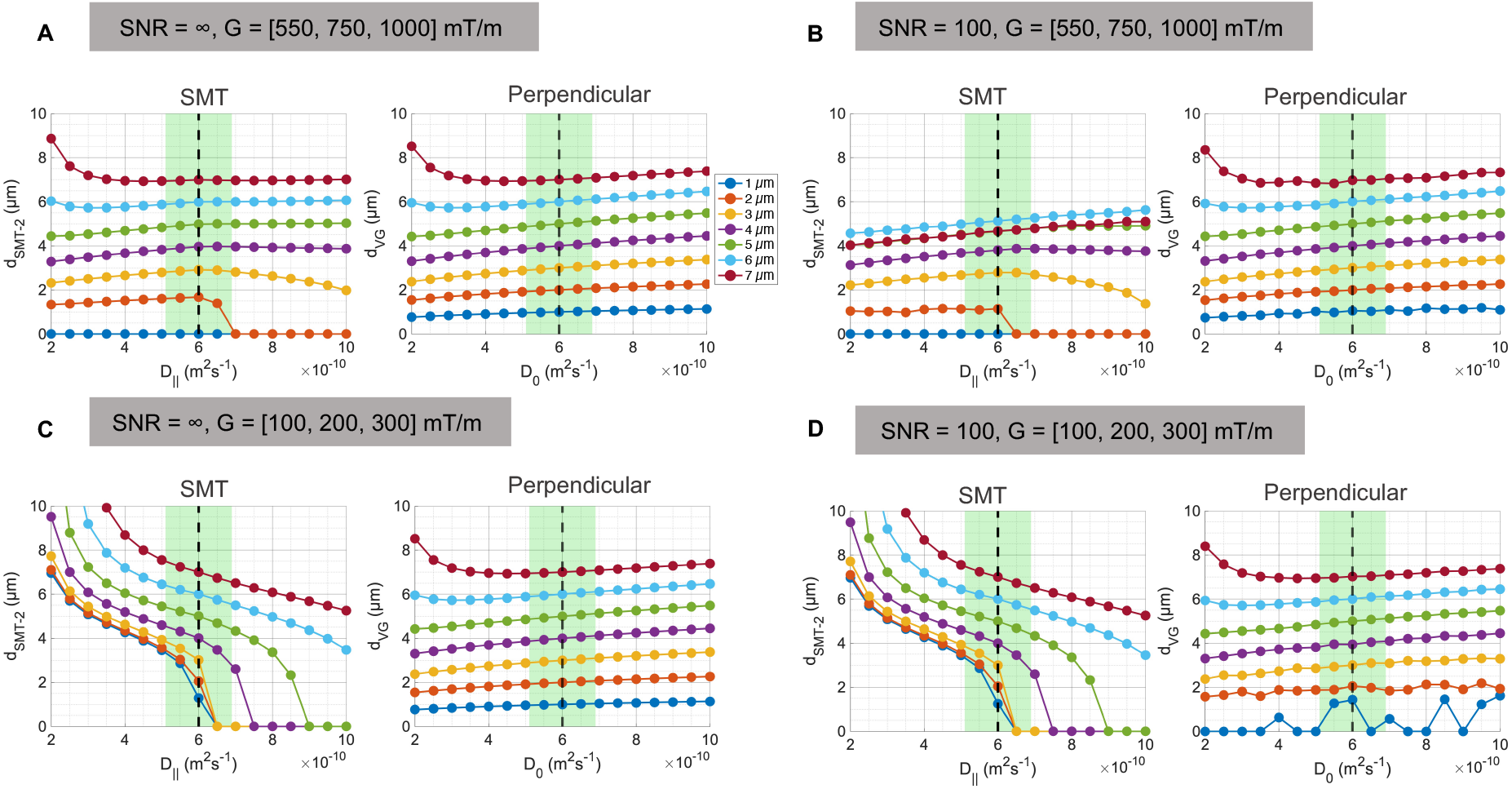
The sensitivity of *d*_SMT-2_ and *d*_VG_ to assumed value of *D*_‖_ depends on *b*-value regime. Estimated *d*_SMT-2_and *d*_VG_ in cylinders vs. assumed value of *D*_‖_ for high diffusion weighting A) *b =* [19.25, 35.79, 63.62] ms *μ*m^-2^ and SNR = ∞, B) b = [19.25, 35.79, 63.62] ms *μ*m^-2^ and SNR = 100 (Rician), and lower diffusion weighting C) b = [0.636, 2.545, 5.726] ms *μ*m^-2^ and SNR = ∞ and D) *b* = [0.636, 2.545, 5.726] ms *μ*m^-2^ and SNR = 100 (Rician). The signal was generated using 30 isotropically distributed directions, *f*_*a*_ = 1 and PGSE parameters *δ* = 7.1 ms and Δ = 20 ms. The higher b shells use *G* = [550, 750, 1000] mT/m and the “low b” shells use *G* = [100, 200, 300] mT/m. The true *D*_‖_ = 0.6 *μ*m^2^ms^-1^, is marked by the black striped line, and *D*_‖_± 15% is represented by the green shaded area. The datapoints represent the mean of *n* = 50 repeats.

Using an incorrect value of *D*_‖_ to within ±15% of the ground truth did not noticeably change *d*_SMT-2_ for cylinders with *d* > 2 *μ*m at high *b*-values and infinite SNR in Fig. 5A. For diameters > 2 pm, there was thus little dependence of *d*_SMT-2_ on the value of *D*_‖_. For cylinders with *d* = 2 *μ*m, the use of a larger-than ground truth *D*_‖_ caused SMT-2 to fail and diameters of 1 *μ*m could not be resolved, regardless of the assumed value of *D*_‖_ The value of *d*_VG_ showed some dependence, albeit small, on the assumed value of D_0_ for diameters > 1 *μ*m. The inclusion of Rician noise in Fig. 5B caused an increased dependency of *d*_SMT-2_ on the assumed value of *D*_‖_, apparent from the increased slope of the plots. As expected from Fig. 3, the Rician noise caused a general underestimation of all diameters, particularly evident in the larger diameters for which the fit misattributed the large signal attenuation to a decreased signal fraction (e.g. 7 *μ*m in Fig. 5B). For the low *b*-values in Figs. 5C-D, there was a much clearer dependence of *d*_SMT-2_ on *D*_‖_. The trends were almost identical for the noise-free/noisy conditions.

### Diameter estimation in segmented axons from XNH volumes of the vervet monkey WM

We found that the fiber architecture, axonal OD and axonal microdispersion differed considerably between the splenium and crossing fiber regions of the vervet monkey brain. The axons segmented from the splenium region, shown in Fig. 6A, were significantly smaller (mean AD = 2.75 *μ*m, SD = 0.53 *μ*m) and exhibited a narrower ADD than those from the crossing fiber region (mean AD = 4.00 *μ*m, SD = 1.26 *μ*m) in Fig. 6B. In comparison to axons from the organised CC environment, the axons from the crossing fiber region were very heterogeneous in terms of length, diameter, shape and OD. The thinnest, thickest and longest axons are shown in Fig. 6D-F. To measure the axial diffusivity, the main directions of the axons were first calculated via a principal component analysis of their trajectories. Intra-axonal diffusion of spins was simulated for up to 100 ms and the displacements of the spins were recorded in the respective main direction of each axon. From the mean-squared-displacements of the spins, the diffusion coefficient in the main direction, here denoted as the z-direction *(D*_*z*_), was approximated and its variation with diffusion time, *t*_*d*_, is shown in Fig. 6G-E for the splenium and crossing fiber axons respectively. The values of *D*_*z*_ in the splenium axons were higher than those in the crossing fiber region, owing to the more irregular trajectories of the crossing fiber axons as seen in Fig. 6F.

**Fig. 6:**
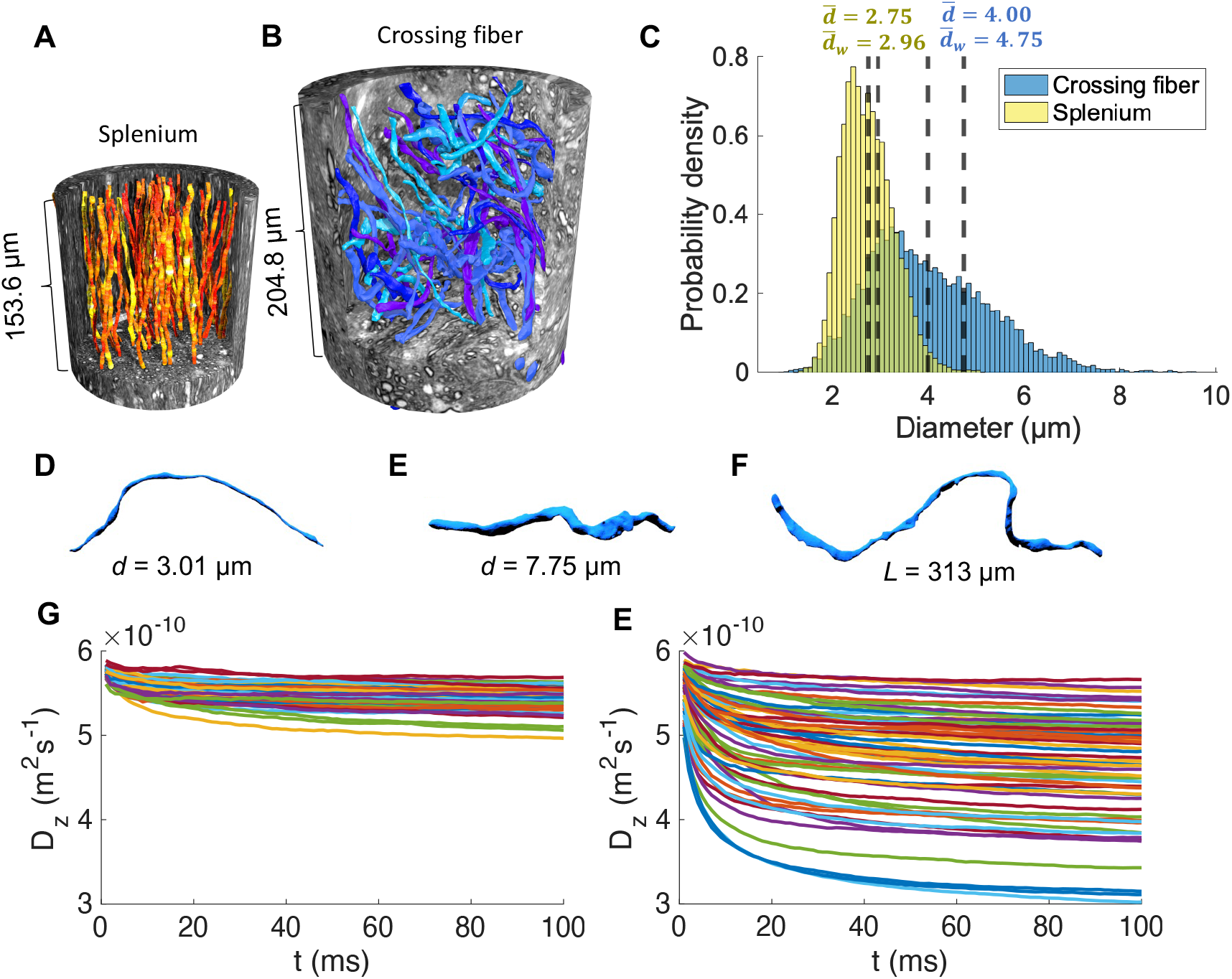
Properties of real axons from different WM fiber architectures in the vervet monkey brain. 3D reconstructions of A) 54 splenium axons (segmented at 75 nm isotropic resolution) and B) 58 crossing fiber axons (segmented at 500 nm isotropic resolution) in their respective XNH volumes. C) Combined 3D AD distributions over all measured diameters in the splenium (yellow) and crossing fiber region (blue). The striped lines mark the means, 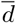, or volume-weighted means 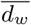, of the distributions. The D) thinnest, E) thickest and F) longest axons from the crossing fiber region demonstrate the significant variability of axonal morphology that can exist on the MRI subvoxel scale. G) and E) show the variation of the diffusion coefficient, *D*_*z*_, in the respective main direction of each axon with diffusion time *t*_*d*_ (data points every 1 ms per axon). Different colours represent different axons.

For evaluation of the SMT and PL implementations in the real IAS and under different conditions, we simulated four acquisitions within the axons from the splenium and crossing fiber regions. To isolate the effects of the real axonal geometries on the estimated diameter, the SNR was set to ∞. The four acquisitions consisted of three shells sampled in 30 gradient directions each due to the similar lower bounds obtained using 30 and 512 directions in Fig. S1. The acquisitions used either a high or low gradient strength set, and either a short (*t*_*d*_ = 12.7 ms) or a long (*t*_*d*_ = 37.7 ms) diffusion time. The intrinsic diffusivity *D*_0_ was set to an ex vivo diffusivity of 0.6 *μ*m^2^ms^-1^. SMT-2, SMT-3 and the PL were fitted to the PA signals to obtain estimates of *d*_SMT-3_, *D*_‖_from the SMT-3 fit, *d*_SMT-2_ and *d*_PL_, as shown in Fig. 7.

**Fig. 7:**
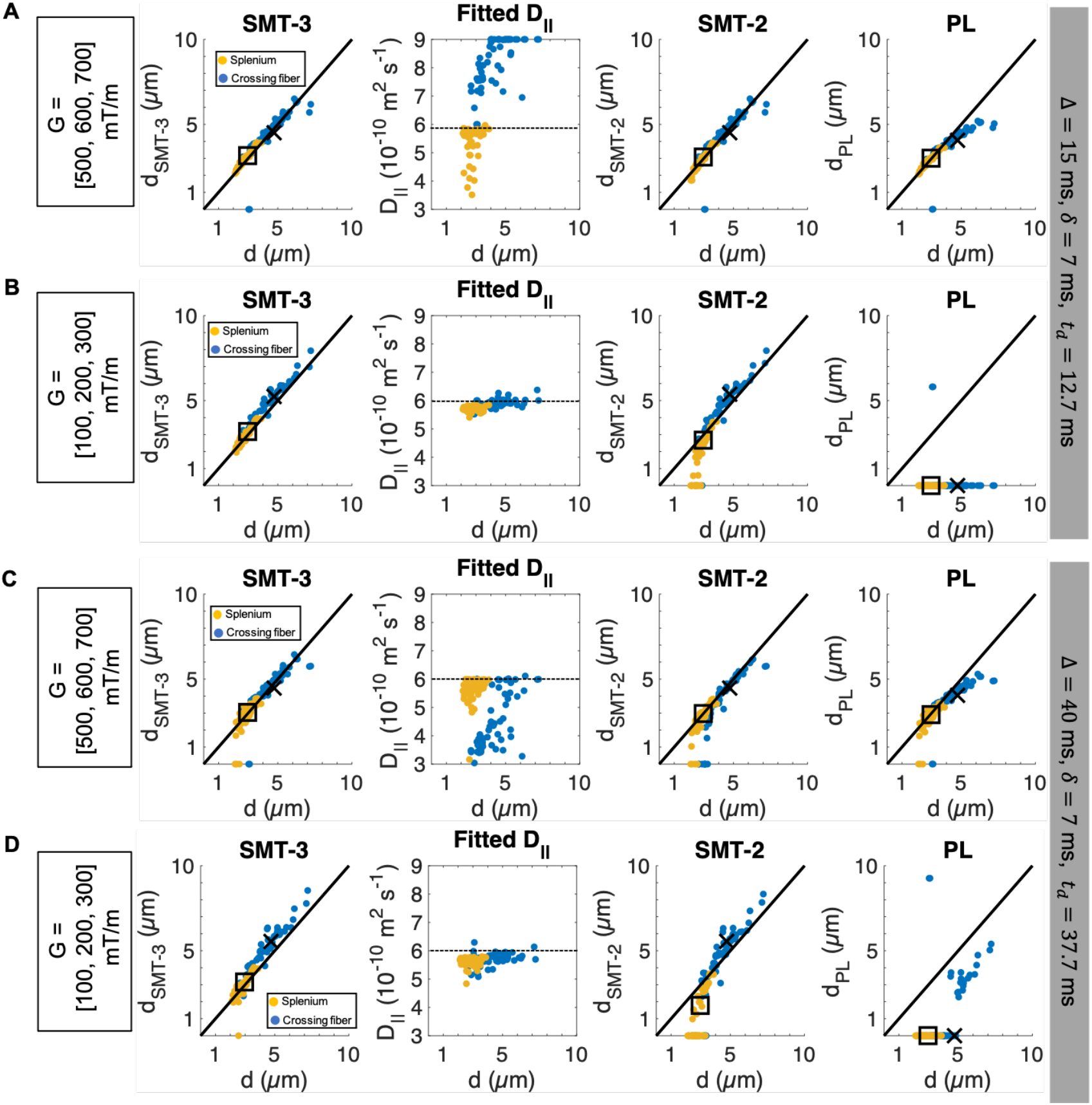
In real axons, a fixed *D*_‖_ can be assumed for high *b*-values. The estimated parameters *d*_SMT-3_, *D*_‖_, *d*_SMT-2_ with assumed ex vivo *D*_‖_ = 0.6 *μ*m^2^ ms^-1^ and *d*_*PL*_ are plotted against the volume-weighted AD of the 54 axons in the splenium (yellow) and the crossing fiber region (blue). The parameters are calculated for different combinations of short (Δ = 15 ms) and long (Δ = 40 ms) gradient separations with high *(G* = [500, 600, 700] mT/m) or low (G = [100, 200, 300] mT/m) gradient strengths, as indicated. This gave three-shell acquisitions with A) *b* = [11.11, 16.00, 21.77] ms *μ*m^-2^, B) *b* = [0.549, 2.198, 4.945] ms *μ*m^-2^, C) *b* = [33.03, 47, 56, 64.73] ms *μ*m^-2^ and D) b = [0.795, 3.180, 7.155] ms pm^-2^. The signals were generated with MC simulations using ex vivo *D*_0_ = 0.6 pm^2^ms^-1^ and were sampled in 30 gradient directions. For all acquisitions, *δ* = 7 ms and SNR = ∞ (barring the intrinsic noise associated with MC simulations). Square marker: volume-weighted AD of splenium axon population, cross marker: volume-weighted AD of crossing fiber population.

At high *b* (Fig. 7A,C), *d*_SMT-3_ and *d*_SMT-2_ provided accurate approximations of both the individual AD and the volume-weighted mean AD of the splenium/crossing fiber axon populations at short and long diffusion times. The estimated *d*_PL_ was accurate for the splenium axons, but underestimated those of the larger crossing fiber axons, in agreement with the breaking down of the Neuman limit for large axons in Fig. S5A. The *D*_‖_ estimates from the SMT-3 were scattered over the range of possible *D*_‖_ values. As shown in Fig. S7, a finite SNR of 100 caused an underestimation of the largest diameters for the high *b*-value acquisition in Fig. 7C, in line with the underestimation of large diameters in Fig. 3.

Using lower *b* and short diffusion times, as in Fig. 7B, *d*_SMT-3_ slightly overestimated the diameters of all axons. There was a subtle positive correlation between *d* and *D*_‖_, the values of which were similar to or higher than *D*_*z*_ in Fig. 6G-H. *d*_SMT-2_ underestimated the diameters of the smaller splenium axons, but overestimated those from the crossing fiber region. The PL fit failed for all but one of the axons, which was greatly overestimated (Fig. 7B). Interestingly, at longer diffusion time, as in Fig. 7D, there was a further overestimation of *d*_SMT-3_ for the crossing fiber axons and a shift in *D*_‖_ towards lower values. In *d*_SMT-2_, the under- and overestimation of the splenium and crossing fiber axons respectively were both enhanced. At this longer diffusion time, *d*_PL_ was non-zero for some of the larger crossing fiber axons, but was significantly underestimated.

To evaluate the methods of AD estimation with in vivo diffusivities and gradient strengths offered by Connectom MRI scaners, the simulations were repeated for *G* = [100, 200, 300] mT/m and two different diffusion times, using an intrinsic diffusivity of *D*_*0*_ = 2 *μ*m^2^ms^-1^. The results are shown in Fig. 8. Strikingly, the volume-weighted mean diameter of the splenium axon population was only non-zero for one metric and under one condition: it was accurate for *d*_*SMT-3*_ at the shortest investigated effective diffusion time *t*_*d*_ = 12.7 ms in Fig. 8A. At the longer diffusion time, *d*_sMT-3_ provided accurate estimates of a subset of the individual axons, but the mean of the population could not be fitted. The individual and population mean ADs of the splenium axons could not be estimated using the SMT-2 or PL approaches for any parameter combinations. The ADs of the crossing fiber axons were generally somewhat overestimated by SMT-3 at all diffusion times. Contrary to the splenium axons, SMT-2 could fit the ADs of the largest crossing fiber axons. The PL implementation could fit very few axons at short diffusion times (Fig. 8A), but its performance improved with increasing diffusion time (Fig. 8B). As in the simulations with ex vivo diffusivities, the values of of *D*_‖_decreased with increasing diffusion time.

**Fig. 8:**
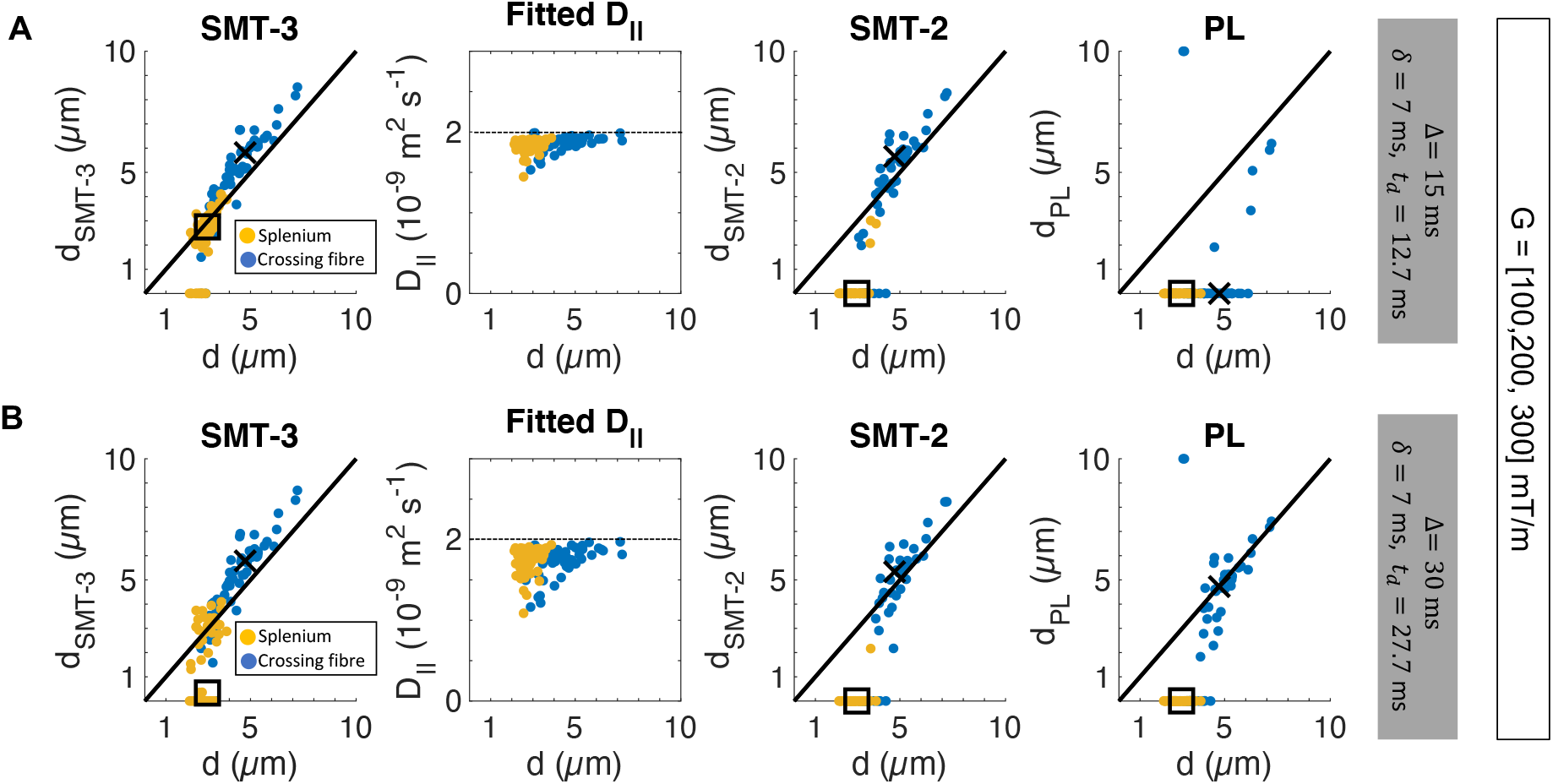
Axon diameter estimation in real axons using an in vivo intrinsic diffusivity, *D*_0_ *=* 2 *μ*m^2^ms^-1^. The estimated parameters *d*_SMT-3_, *D*_‖_, *d*_SMT-2_ and *d*_PL_ are plotted against the volume-weighted AD of the 54 axons in the splenium (yellow) and the crossing fiber region (blue). The parameters are calculated for acquisitions with A) Δ = 15 ms and *b* = [0.44, 1.78, 4.00] ms *μ*m^-2^ and B) Δ = 30 ms and *b* = [0.97, 3.88, 8.73] ms *μ*m^-2^. Each acquisition consisted of three shells with 30 gradient directions, *G* = [100, 200, 300] mT/m, *δ* = 7 ms and SNR = ∞. Square marker: volume-weighted AD of splenium axon population, cross marker: volume-weighted AD of crossing fiber population

## Discussion

By segmenting axons from synchrotron X-ray nano-nolotomography volumes of a splenium and crossing fiber region of the vervet monkey brain, we explored the impact of real axonal morphologies on axon diameter estimation with powder average diffusion MRI approaches. Simulations of diffusion within the segmented axons, using both in vivo and ex vivo diffusivities, show that accurate measures of mean axon diameter can be obtained. In the real axonal substrates, we also find that the diameter estimates exhibit a time dependence, provided that the *q*-values of the applied sequence are low enough to preserve the diffusion MRI signal arising from large displacements. Given the broad distribution of axon diameters found in the crossing fiber region, it is imperative that the diffusion MRI acquisition has sensitivity to both small (including those too small to be segmented from the XNH volumes) and very large diameters. We show here how powder average-based methods are subject to upper and lower bounds of measurable diameter. Although these depend on the gradient strength [14, 56, 57], they also depend on the *b*-value and the number of gradient directions. Ultimately, we find that the effective SNR of the measurement is the key limiting variable of measurable diameter, and we show the importance of removing Rician bias from the diffusion MRI signal.

### Sequence parameter considerations

#### The effect of *q*-value, *b*-value and the number of gradient directions on the bounds of measurable diameter

We show that sensitivity profiles of PA-based methods to different diameters (Fig. 2B) are influenced by sensitivity of the acquisition to different angles within the cylindrical micro-domains (Fig. 2C). The *q*-value and the diffusion time of the acquisition act as spatial filters, restricting the maximum detectable displacements of the spins. A higher *q*-value increases the sensitivity of the acquisition to smaller length scales in directions perpendicular to the cylinder. However, we show that if the *q*-value is high enough to cause attenuation of the signal from an ensemble of spins before it has diffused for the entire diffusion time, the acquisition loses sensitivity in the axial direction. For a given diffusion time and SNR, we show that an increase in *q* narrows the angular sensitivity profile. Although an increased gradient strength will move the lower bound to smaller diameters, in accordance with the findings of Dyrby et al. [14] and Sepehrband et al. [56], the upper bound will also be reduced, narrowing the range of measurable diameters. This is true also for methods that estimate AD from measurements perpendicular to axons [10, 11, 13–16], but the increased attenuation of the PA due to its averaging across many directions and length scales typically incurs a higher location of the lower bound, as shown in the comparison between *d*_SMT-1_ and *d*_VG_ in Fig. S1.

We found that the number of uniformly distributed gradient directions influenced the lower bound of measurable AD, with smaller diameters demanding measurements with higher angular resolution. This is in line with the findings of Li et al. [58] who show that at SNR= ∞, the number of directions determines how accurately the measured PA signal reflects the ground truth signal. The angular resolution thus places a lower bound on the measurable AD, separate to that incurred by the sequence parameters and finite SNR. We demonstrate that with Rician noise, increasing the angular resolution cannot decrease the SNR-incurred lower bound. For Gaussian noise, however, the higher number of sampling points given by the higher angular resolution may increase the effective SNR and provide access to smaller diameters. In practice, it has been shown that it is better to increase the number of directions than to perform many repeats of the same shell [59]. Furthermore, an increased angular resolution increases the robustness of the AD estimate to different underlying fiber configurations and OD, as seen in Fig. S2. This agrees with other studies that find that more gradient directions lower the variance in parameter estimates from the PA signal [33, 60]. Our results also indicate that an increased angular resolution does not yield noticeably increased rotational invariance after a certain number of directions, similar to the findings of [61] where little improvement in the rotational invariance of the fractional anisotropy measurement was found beyond 20 directions at *b* =1 ms *μ*m^-2^.

All simulations on cylinders in this investigation assume a single cylinder direction. In practise, this is not realistic even in the CC [19, 20]. For methods that assume a single fiber direction [10, 11, 13, 14], this is a limitation. PA methods, on the other hand, may benefit from fiber dispersion as it has been shown that the less anisotropy there is on the voxel scale, the fewer directions are required to obtain rotational invariance of the PA [58, 60, 61]. It is thus possible that axon diameter estimates in ordered WM regions such as the CC require the use of more gradient directions than in heterogeneous crossing fiber regions.

#### Is it necessary to fit the intra-axonal parallel diffusivity?

We show here that the need to fit *D*_‖_depends on the sequence parameters used. For high *b*-values, the signals at low *α* (roughly parallel to the cylinder axis) were heavily attenuated and contributed little to the overall PA. The dependency of the estimated diameter on *D*_‖_ was therefore low and the assumption of erroneous *D*_‖_and *D*_*0*_ to within ±15% of the true value did not have a significant impact on the estimated diameter (Fig. 5A-B). At lower *b*, as in Fig. 5C-D, on the other hand, the contribution of the low *α* signals to the overall PA was more significant and the dependency on *D*_‖_ could not be ignored. Some AD estimation methods, such as ActiveAx [13] and the SMT implementation by Fan et al. [34] assume a value of Dp Other methods e.g. AxCaliber [10, 11] and the PL implementation [35] (either directly or indirectly) fit it. For PA methods, this entails that estimates of *D*_‖_ calculated by measuring the signal parallel to axons in the CC are likely to be sufficient at ex vivo intrinsic diffusivities for b ≥ 20 ms *μ*m^2^. For methods that implement lower *b*-values, *D*_‖_ may need to be fitted (in addition to the signal from other tissue compartments), especially considering the time-dependence of the intra-axonal axial diffusivity [19, 62–64].

#### Choice, but not number, of *b*-values affects measurable range of diameter

Fitting the SMT and PL implementations to the signal from different sets of *b*-values (all b ≳ 20 ms *μ*m^2^ to simulate suppression of the E AS signal [35]) showed that there may be little advantage to increasing the *q*–value (and thus the *b*-value) of an acquisition. This supports the trend in Fig. 2B, in which an increase in *q*-value after a certain point does little to lower the lower bound, but narrows the range of measurable diameter. Furthermore, the finding of no noticeable advantage of densely sampling many *b*-values lends support to the optimisation framework of Alexander et al. [65] in which the optimal acquisition design for axon diameter estimation used few shells (3 to 6), and to the approach of ActiveAx [13]. Our results thus indicate that it may not be necessary to perform a dense sampling of *b*-values as in the SMT implementation of Fan et al. [34], AxCaliber [11] and the PL implementation [35]. A similar suggestion was recently made by Veraart et al. [66]. Moreover, while the sensitivity criteria of Nilsson et al. [43] provided indications of the range of measurable diameter for SMT-1 fits to a single *b*–value (Fig. S1, Fig. S3), no equivalent metric exists for a multi-shell fit, making simulations of the signal important in predicting the sensitivity of a multi-shell acquisition to diameter.

The drops to zero at small and large diameters resulting from multi-shell fits for Rician distributed noise in Fig. 3 introduce a problematic degeneracy: axons with diameters above the upper bound of measurable diameter may instead appear as smaller axons. We show here the existence of axons up to 8 *μ*m in mean diameter in the crossing fiber region. In the Rician regime and high *b*-values, as in Fig. 3, these are underestimated by both the SMT and PL approaches. Therefore, to work in the Rician regime one needs to ensure that the upper bound is sufficiently high to prevent the largest axons in the tissue from being underestimated. Sensitivity to larger axons may be gained by using longer diffusion times [14], and -as we show in Fig. 2B -by increasing the effective SNR or reducing the *q*-value. We show here that the use of Gaussian distributed noise also ensures improved sensitivity to larger diameters for both the SMT and PL approaches. As in the results of Fan et al. [34], we observed a relationship between estimated diameter and IAS signal fraction at large diameters for the SMT-based approach. This mostly disappeared at infinite SNR, indicating that it may not be an issue at lower *b*-values, where the signal attenuation is lower and effective SNR is higher. Our analysis also revealed that the apparent underestimation of diameter in *d*_PL_ compared to *d*_SMT-2_ in Figs. 3 and 4 is due to large axons falling outside of the assumed Neuman limit (Eq. 5) which was used to calculate a diameter from *D*_⊥_ in the PL implementation (the SMT implementations instead used the full expression in Eq. 4).

#### The importance of the SNR and noise type

For a single *b*-value, we show here how an increase in SNR (assuming Gaussian distributed noise) increased the angular sensitivity range of the acquisition (Fig. 2), as expected from the sensitivity criteria presented in [43]. Similarly, for fits of diameter to multiple high *b*-value signal sets, we found that Gaussian distributed noise resulted in a wider range of measurable diameters (Fig. 3) and provided more accurate estimates of the intra-axonal signal fraction. As such, Gaussian distributed noise also prevented the systematic underestimation of diameter that is seen with Rician noise. This agrees with the findings of Fan et al. [34], in which it is also argued that the use of real-valued diffusion MRI data with Gaussian noise is more independent of the underlying fiber orientation distributions. The advantages of using Gaussian distributed noise stem from the fact that powder averaging of signals with a noise distribution centred around the mean avoids bias in the estimated average of the signal. In light of this, recovery of the real-valued diffusion MRI data with Gaussian distributed noise by performing a phase correction [67] is likely an important step in improving both the range and accuracy of measurable ADs.

#### Microdispersion affects PA-based axon diameter estimates in segmented axons from the vervet monkey brain

At high *b*-values and ex vivo intrinsic diffusivities (Figs. 7A and C), we show that wide range of ADs from the splenium and crossing fiber WM regions could be accurately estimated using the SMT implementation, regardless of diffusion time. The PL underestimated some of the larger crossing fiber axons, in agreement with the breaking down of the Neuman limit for large axons that we observe in Fig. S5A. Our simulations in real axons thus suggest that, in practice, the assumption of the Neuman limit cannot necessarily be made for all WM voxels and acquisitions; in our data, it was an adequate assumption for the splenium CC region, but not the crossing fibre region. Hence, even if the acquired diffusion MRI signals and calculated *D* _⊥_ hold the anatomical information to provide an accurate AD estimate, the method by which a diameter is calculated from *D* _⊥_ may bias the estimate. The formulation by van Gelderen et al. [47] (Eq. 4) generalises across sequence parameters and diameters and may thus provide a more optimal sensitivity profile to a broad range of ADs for given acquisition parameters. Furthermore, the scattered and inaccurate estimates of *D*_‖_ from SMT-3 (some even surpassing the intrinsic diffusivity in Fig. 7A), yet the accurate *d*_SMT-3_ estimates of AD, support our finding that the estimated AD is not very sensitive to inaccuracies in *D*_‖_ at high *b*-values (Figs. 5A and B). Unlike Lee et al. [22], we did not observe an overestimation of AD.

At lower *b*-values, the effects of axonal microdispersion manifested as a slight overestimation of the crossing fiber axon ADs that increased with increasing diffusion time. Fitting *D*_‖_ was necessary to obtain accurate AD estimates for the smaller splenium axons. Despite the extremely tortuous trajectories of the crossing fiber axons, their fitted *D*_‖_ were generally higher than those of the smaller splenium axons. It is possible that axons exhibit similar trajectories somewhat independently of the voxel-scale fiber architecture, given that these are modulated by obstacles in the local environment [19]. If so, the correlation between *D*_‖_ and diameter could be due to spins in smaller axons probing the curvature of the IAS to a greater extent. The curvature could be caused by microdispersion and diameter variations [19, 22, 63, 68]. In contrast to the PA estimates of *D*_‖_, the diffusivities measured in the single main direction of each axon *(D*_*z*_) were markedly lower in the crossing fiber region than in the splenium (Figs. 6G and E). Lastly, we observed a decrease in *D*_‖_ at longer diffusion times, agreeing with the time-dependence of *D*_*z*_ in Fig. 6. The nature of this time dependence could be indicative of the diameter variations and degree of microdispersion of the axons [19, 63], and thus also an indication of the density of cells or other extra-axonal structures in the WM [19]. The time dependence of both *D*_‖_ and the diameter could potentially act as biomarkers of situations where the WM cell density is expected to change, such as in pathology or inflammation.

The simulations within the realistic IAS at in vivo diffusivities (Fig. 8) highlight the importance of mapping the sensitivity profile of the whole acquisition setup including the model and acquisition parameters. Even with gradient strengths accessible to human diffusion MRI experiments only via Connectom scanners, the splenium ADs could be accurately estimated only by SMT-3 at the shortest diffusion time and provided that *D*_‖_ was fitted. At increasing diffusion times, the volume-weighted mean AD of the splenium population could not be estimated. Additionally, the overestimation of the ADs of the crosssing fiber axons at all diffusion times could be explained by the high in vivo intrinsic diffusivity entailing that spins probe larger distances, and thus more microdispersion, than at ex vivo intrinsic diffusivities. The lack of time-dependence in the crossing fiber ADs could be due to the spatial filtering effect of the *q*-value.

### Limitations

This investigation restricted the analysis of PA-based AD estimates to the IAS, for different sequence parameters and SNRs. At high *b*-values, the observation of a signal decay proportional to *b*^-0.5^ indicates that the signal mostly arises from thin, cylindrical structures [39, 40, 69], and that the EAS is suppressed. However, in the splenium XNH volume, we observed cell clusters and vacuoles that together constituted 6.1% of the total volume fraction [19]. Recent studies show that the cell somas could contribute to the PA signal at short diffusion times [70], complicating the SMT and PL fit to the PA signals. The presence of any restricted or hindered compartment from which the signal remains at high *b*–values will complicate the fits, unless it is explicitly modelled. These compartments could include e.g. irregularities in the axonal myelin or cellular processes. The observed dot compartment in ex vivo tissue [13, 35] -completely restricted in all directions -will systematically bias PA-based AD measurements, although its contribution to the signal has been shown to be negligible in vivo [40, 71, 72].

Furthermore, use of the single-compartment PL implementation as in Veraart et al. [35] requires high *b*-values both for the suppression of the EAS and to fulfil the assumptions of the PL model. In Fan et al. [38] and Veraart et al. [35], the use of Connectom scanners for the in vivo applications enabled high gradient strengths, and thus high *q* and *b*-values. On regular clinical scanners with limited gradient strengths, high *b*-values could be achieved with longer diffusion times, but this would be at the cost of a reduced sensitivity to small diameters and a long echo time that would reduce the SNR of the acquisition. Longer effective diffusion times could alternatively be obtained via stimulated echo acquisitions [73, 74], but also at the cost of lower SNR.

At lower *b*-values, the PL is no longer valid and only the SMT implementation provides an adequate fit to the PA signals. Given that the EAS is not suppressed at such *b*–values, the signal contributions from other compartments must be modelled, as in Fan et al. [34]. The accuracy of the AD estimate thus depends not only on how accurately the geometry of other compartments are modelled, but also the compartmental *T*_2_ relaxation times [75] and potential exchange rates. However, the sensitivity to microdispersion and *D*_‖_ at low *b*-values is higher, and the time-dependence of *d*_SMT-3_ and *D*_‖_ could provide valuable insight into axonal morphology [20, 22, 41, 43, 63, 76].

One key challenge to AD estimation with diffusion MRI is that real WM voxels contain an ADD, and not single diameters. Here, we have segmented the upper tail of the ADD. Diffusion MRI-based estimates of mean diameter are heavily weighted by the tail of the ADD [35, 77] and the larger axons -like those presented in this study -thus significantly contribute to the signal. In alignment with Dyrby et al. [9, 14] we show that the sequence parameters and the PA model together result in an AD sensitivity profile, with an upper and lower bound of detectable diameter. Despite the heavier weighting of larger axons, axons below the lower bound of measurable diameter (or above the upper bound) still contribute to the total signal and will cause an underestimation of the average AD index, as seen for the splenium axons at in vivo diffusivities.

## Conclusion

We demonstrate that powder averaging techniques can succeed in providing accurate estimates of axon diameter, even in a complex crossing fiber region of the vervet monkey brain. To succeed, the acquisition must have a broad sensitivity profile to different length scales. This is important partly due to the many different axon sizes present within a voxel, as presented here, but also because the powder average probes different length scales in anisotropic micro-domains. Furthermore, we show how the gradient strength, diffusion time and number of gradient directions, as well as the SNR and type of noise distribution, influence the lower and upper bounds of measurable diameter. Finally, at low *b*-values we show that the acquisition becomes sensitive to axonal microdispersion, which could be an interesting biomarker of WM health and pathology. We foresee that this characterisation of the limits and potential of PA-based approaches to AD estimation will contribute to the development of new methods and models to study the WM microstructure with diffusion MRI.

## Acknowledgements

MA, HMK were supported by the Capital Region of Denmark Research Foundation (grant number: A5657) (PI:TD). This work has received funding from the European Union’s Horizon 2020 research and innovation programme under the Marie Skłodowska-Curie grant agreement No. 754462 (MP). HL has received funding from the European Research Council (ERC) under the European Union’s Horizon 2020 research and innovation programme (grant agreement No 804746).

## Notes

### Competing Interest Statement

The authors have declared no competing interest.

