## Supplementary Materials for "Does powder averaging remove dispersion bias in diffusion MRI diameter estimates within real 3D axonal architectures?"

### S1 Effect of number of gradient directions and SNR on estimated diameter

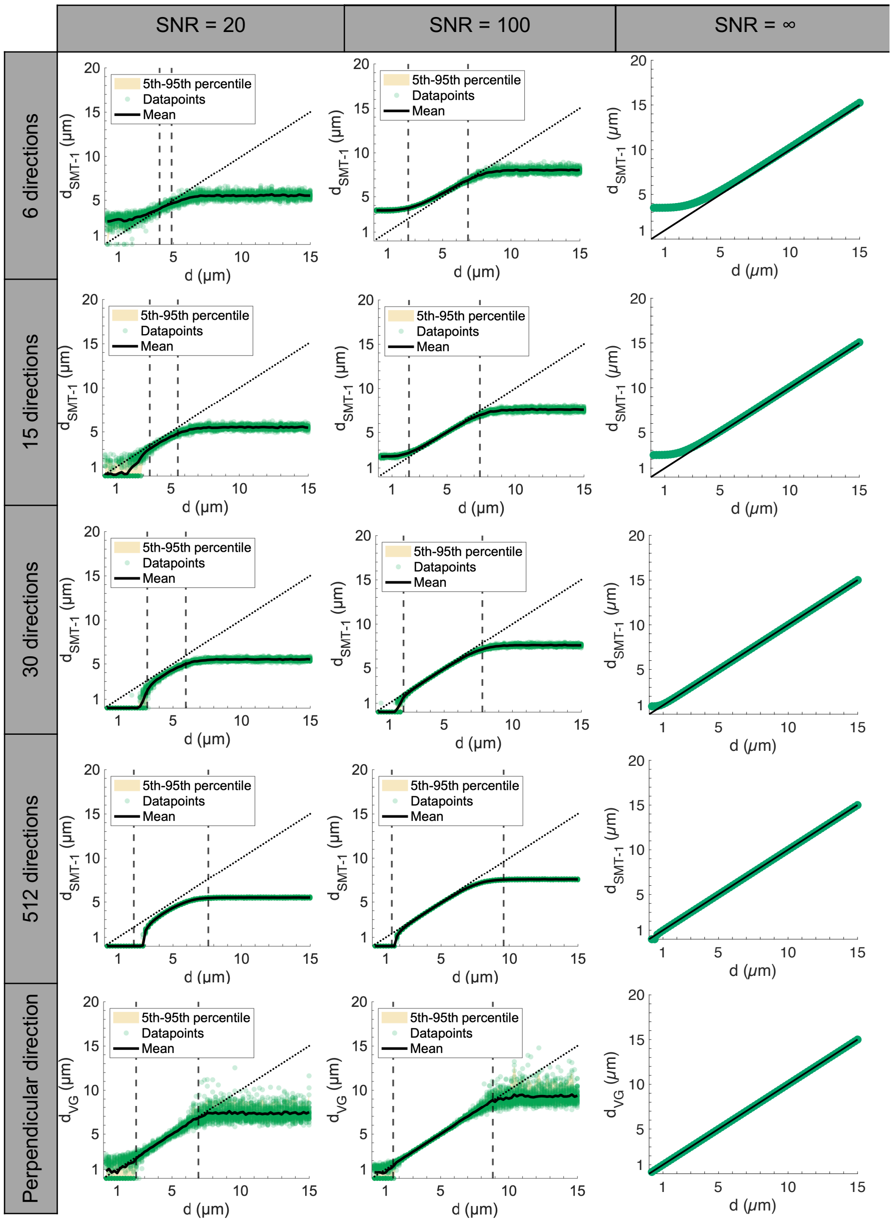


Fig S. 1: **Influence of number of directions and SNR (Rician noise) on the single-shell SMT-estimated diameter, $d_{\mathbf{SMT\text{-}1}}$, in cylinders of diameter between 0.2 and 15.0 $\mu$m at 0.2 $\mu$m intervals.** The single-shell SMT assumes known $f_a = 1$, $D_{\parallel} = 0.6 \cdot 10^{-9}\mathrm{m}^2\mathrm{s}^{-1}$. The lower and upper bounds of measurable diameter are sensitive to both SNR and the number of directions and calc. The last row shows the influence of SNR on the diameter calculated from a single perpendicular direction, $d_{\mathrm{VG}}$, as calculated using Eq. 4 in the main text. $n = 50$ repeats of the acquisition were performed for each diameter, SNR and number of directions. The black striped lines represent the predicted lower and upper bounds of measurable diameter.

The behaviour of $d_{\mathrm{SMT\text{-}1}}$ at small diameters varied when few directions were used (Fig. S1). It either plateaued (as it did using 6 directions) or dropped to 0 (as it did using 15 directions and SNR = 20). This indicated a dependence of the PA signal on the orientation of the gradients relative to the cylinder axis. The sensitivity of $d_{\text{SMT-1}}$ to the orientation of the gradient directions was examined by generating the signals of cylinders that were rotated in 90 isotropically distributed directions. Fig. S2 shows the standard deviation and mean of $d_{\text{SMT-1}}$ estimated from these 90 orientations. In general, the estimated diameter was more rotationally invariant (the standard deviation was lower) the higher the angular resolution.

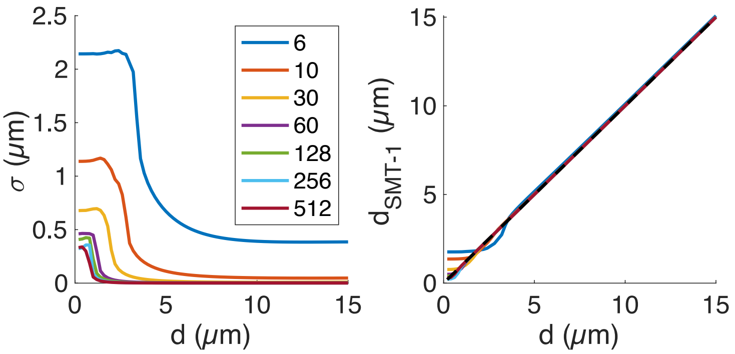


Fig S. 2: **The PA is sensitive to fiber direction.** The standard deviation of $d_{\text{SMT-1}}$ of cylinders rotated around the $y$-axis in 90 uniformly distributed directions, and the estimated mean diameter $d_{\text{SMT-1}}$, depend on the number of gradient directions. PGSE parameters $\delta = 7.1$ ms, $\Delta = 20$ ms and $G = 600$ mT/m were used.

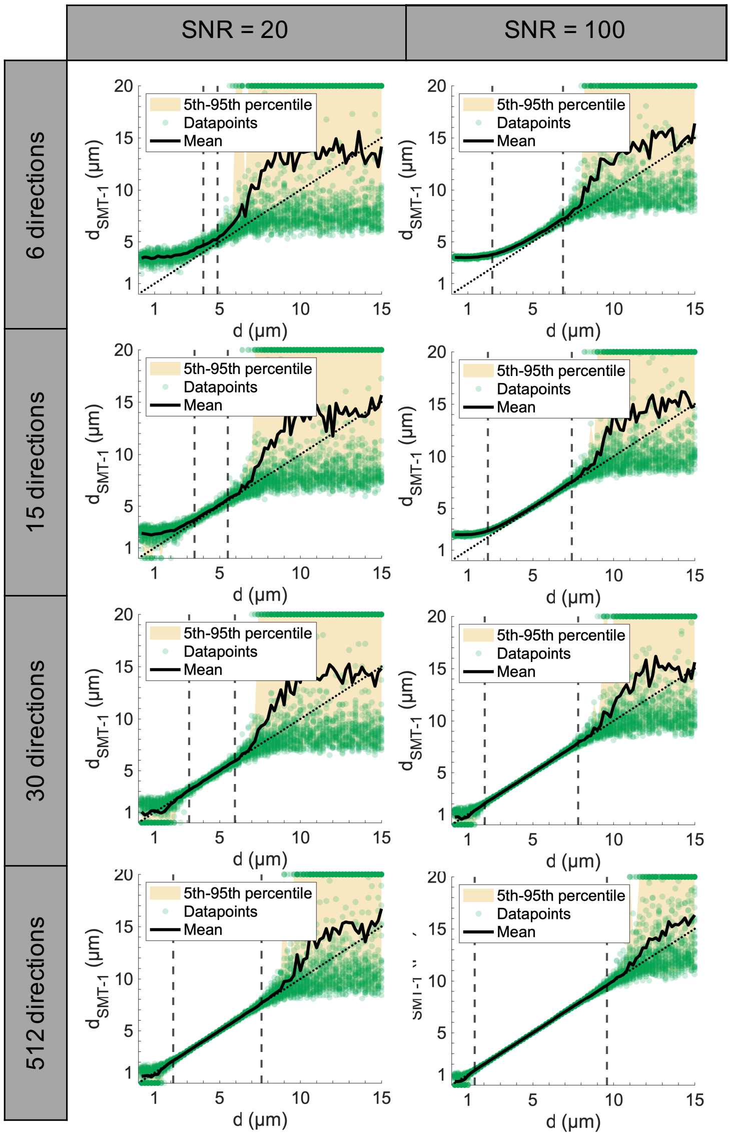


Fig S. 3: **Influence of Gaussian noise and number of directions and on the single-shell SMT-estimated diameter, $d_{\mathbf{SMT\text{-}1}}$, in cylinders of diameter between 0.2 and 15.0 $\mu$m.** The single-shell SMT assumes known $f_a = 1, D_{\|} = 0.6\ \mu\text{m}^2\text{ms}^{-1}$. The last row shows the influence of SNR on the diameter calculated from a single perpendicular direction, $d_{\text{VG}}$, as calculated using Eq. 4 in the main text. $n = 50$ repeats of the acquisition were performed for each diameter, SNR and number of directions. The black striped lines represent the predicted lower and upper bounds of measurable diameter.

#### Other Supplementary Figures

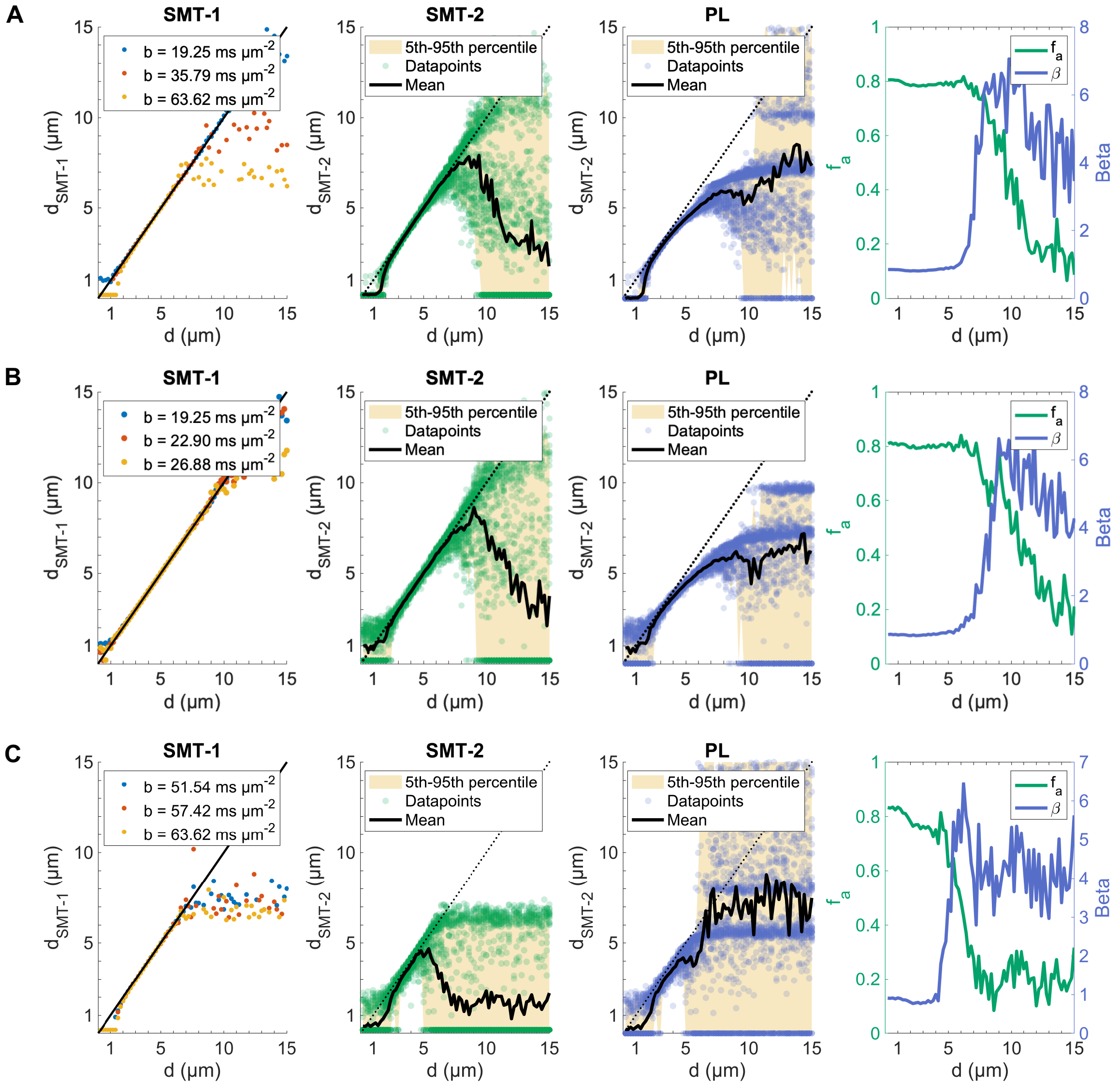


Fig S. 4: **Multi-shell fits with Gaussian noise**. Multi-shell fit of SMT and PL to the signal from cylinders of diameter between 0.2 and 15.0 $\mu$m at SNR = 100 (Gaussian noise) and $f_a = 0.8$. Fitted $d_{\text{SMT-1}}$, $d_{\text{SMT-2}}$, $d_{\text{PL}}$, $f_a$ and $\beta$ for ground truth A) three shells with $b = [19.25, 35.79, 63.62]$ B) three shells with $b = [19.25, 22.90, 26.88]$ ms $\mu$m$^{-2}$ and C )three shells with $b = [51.54, 57.42, 63.62]$ ms $\mu$m$^{-2}$. The signal is generated using $D_{\|} = 0.6$ $\mu$m$^2$ms$^{-1}$, 30 directions, PGSE parameters $\delta = 7.1$ ms, $\Delta = 20$ ms and varying $G$. $n = 50$ repeats of each acquisition were performed for each diameter.

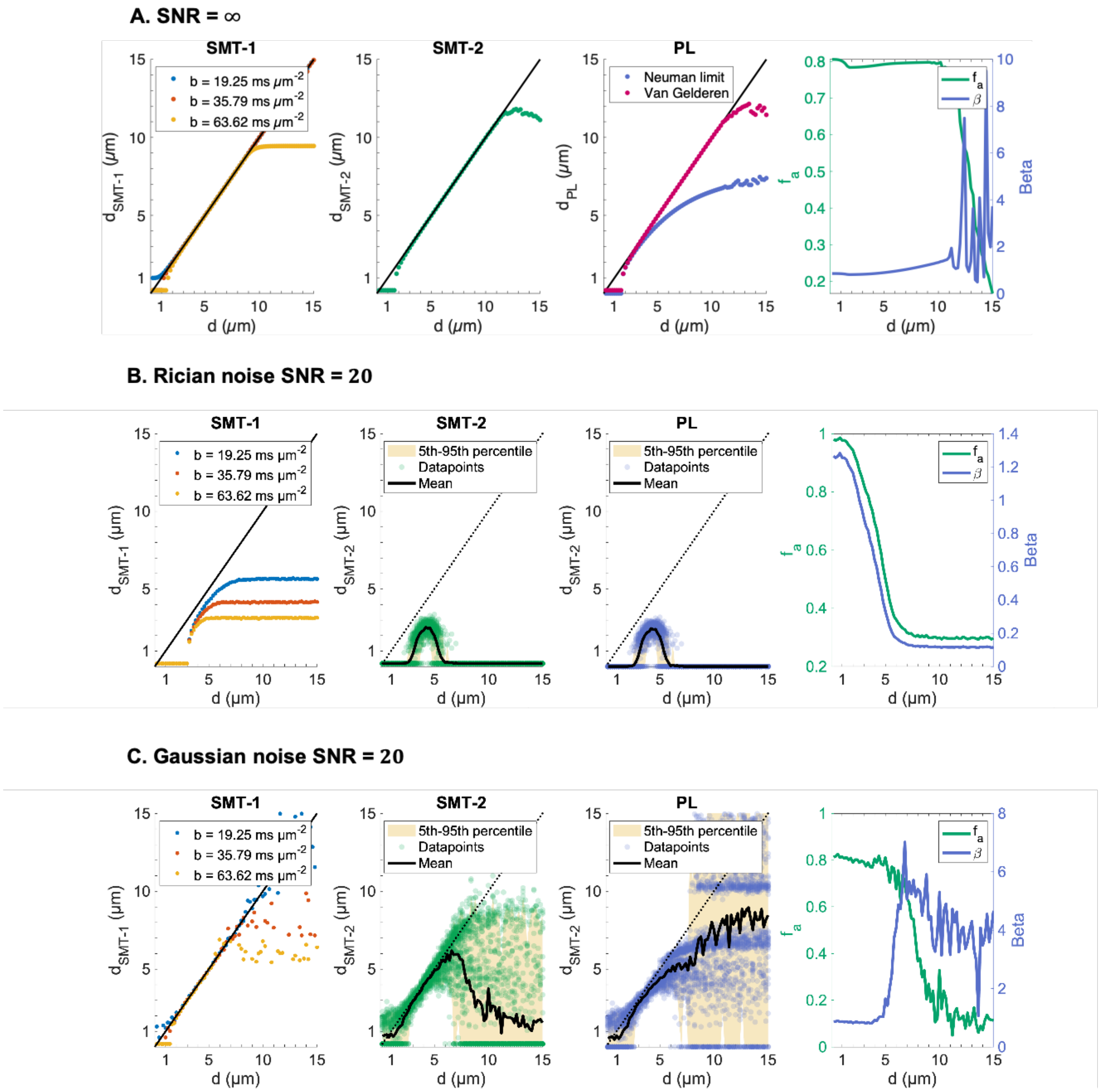


Fig S. 5: **Multi-shell fit at A) SNR $= \infty$, B) SNR $= 20$ with Rician noise and C) SNR $= 20$ with Gaussian noise .** Fitted $d_{\text{SMT-1}}$, $d_{\text{SMT-2}}$, $d_{\text{PL}}$, $f_a$ and $\beta$ for $b = [19.25, 35.79, 63.62]$ ms $\mu\text{m}^{-2}$ in cylinders of varying diameter. In A), $d_{\text{PL}}$ is calculated in two ways: one using the assumption of the Neuman limit in Eq. 5 and the second using the full formulation of the signal perpendicular to a cylinder in Eq. 4 of the main text. The signal is generated using $D_{\|} = 0.6\ \mu\text{m}^2\text{ms}^{-1}$, 30 directions and PGSE parameters $\delta = 7.1$ ms, $\Delta = 20$ ms and $G = [550, 750, 1000]$ mT/m. $n = 50$ repeats of each acquisition were performed for each diameter.

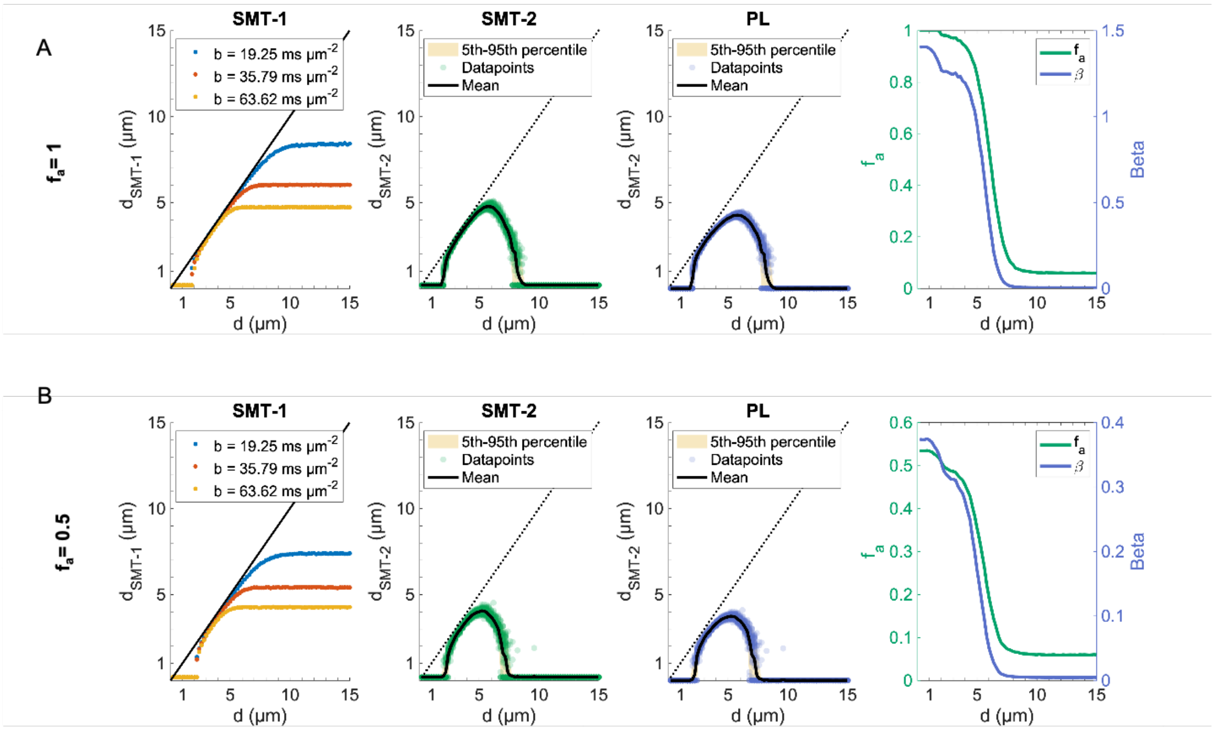


Fig S. 6: **Multi-shell fit of SMT and PL to the signal from cylinders at SNR = 100 (Rician noise) and different $f_a$.** Fitted $d_{\text{SMT-1}}$, $d_{\text{SMT-2}}$, $d_{\text{PL}}$, $f_a$ and $\beta$ for ground truth A) $f_a = 1$ B) $f_a = 0.8$ and C ) $f_a = 0.5$. The signal is generated using $D_{\parallel} = 0.6\ \mu\text{m}^2\text{ms}^{-1}$, 30 isotropically distribued directions and PGSE parameters $\delta = 7.1$ ms, $\Delta = 20$ ms , $G = [550, 600, 650]$ mT/m, $b = [19.25, 22.90, 26.88]$ ms $\mu\text{m}^{-2}$. $n = 50$ repeats of each acquisition were performed for each diameter.

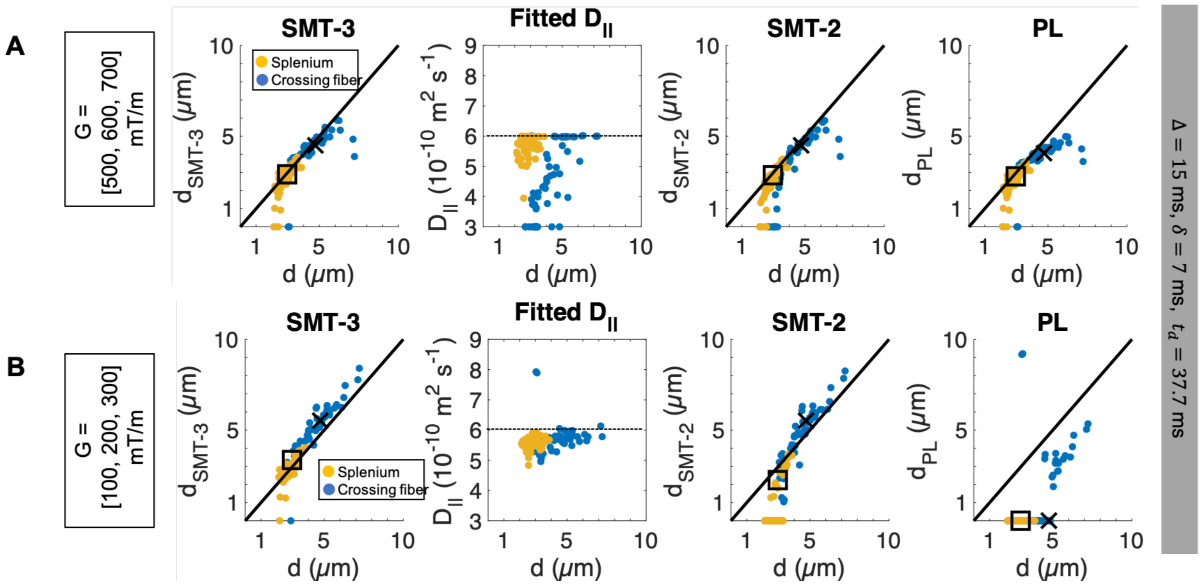


Fig S. 7: **Axon diameter estimation in real axons at finite SNR = 100 (Rician noise)**. The estimated parameters $d_{\text{SMT-3}}$, $D_{\|}$, $d_{\text{SMT-2}}$ and $d_{\text{PL}}$ are plotted against the volume-weighted AD of the 54 axons in the splenium (yellow) and the 58 axons in the crossing fiber region (blue). The parameters are calculated for A) heavy diffusion weighting $b = [11.11, 16.00, 21.77]$ ms $\mu$m$^{-2}$ and B) weaker diffusion weighting $b = [0.549, 2.198, 4.945]$ ms $\mu$m$^{-2}$. For the heavy diffusion weighting, $G = [500, 600, 700]$ mT/m, and for the lower diffusion weighting, $G = [100, 200, 300]$ mT/m, as indicated. For all acquisitions, $\delta = 7$ ms, $\Delta = 40$ ms and SNR = 100. An ex vivo diffusivity of $D_0 = 0.6$ $\mu$m$^2$ms$^{-1}$ was used for the simulations. Square marker: volume-weighted AD of splenium axon population, cross marker: volume-weighted AD of crossing fiber population.
